# SpaceBF: Spatial coexpression analysis using Bayesian Fused approaches in spatial omics datasets

**DOI:** 10.1101/2025.03.29.646124

**Authors:** Souvik Seal, Brian Neelon

## Abstract

Advances in spatial omics enable measurement of genes (spatial transcriptomics) and peptides, lipids, or N-glycans (mass spectrometry imaging) across thousands of locations within a tissue. While detecting spatially variable molecules is a well-studied problem, robust methods for identifying *spatially varying co-expression* between molecule pairs remain limited. We introduce SpaceBF, a Bayesian fused modeling framework that estimates co-expression at both local (location-specific) and global (tissue-wide) levels. SpaceBF enforces spatial smoothness via a fused horseshoe prior on the edges of a predefined spatial adjacency graph, allowing large, edge-specific differences to escape shrinkage while preserving overall structure. In extensive simulations, SpaceBF achieves higher specificity and power than commonly used methods that leverage geospatial metrics, including bivariate Moran’s *I* and Lee’s *L*. We also benchmark the proposed prior against standard alternatives, such as intrinsic conditional autoregressive (ICAR) and Matérn priors. Applied to spatial transcriptomics and proteomics datasets, SpaceBF reveals cancer-relevant molecular interactions and patterns of cell–cell communication (e.g., ligand–receptor signaling), demonstrating its utility for principled, uncertainty-aware co-expression analysis of spatial omics data.

## 1 Introduction

Technological advances in spatial omics^1–3^ have enabled *in situ* profiling of varying molecules, including genes (via spatial transcriptomics (ST))^4–7^, lipids or peptides (using mass spectrometry imaging (MSI))^8–11^, and immune proteins (through multiplex immunofluorescence (mIF))^12–15^, within tissues. The technologies offer distinct yet complementary biological insights, differing in spatial resolution and the number of detectable molecules (throughput). For example, the next-generation sequencing (NGS)-based ST platform Visium (from 10X Genomics)^16^ offers transcriptome-wide gene-expression profiling (throughput ∼ 20, 000) at a 55 *µm* spot-level resolution. MALDI MSI-based platforms (from Bruker Daltonics^17^ and others) offer profiling different types of molecules, such as peptides, lipids, nucleotides, proteins, metabolites, and N-glycans, (throughput ∼ 50 − 1000) at 10 *µm* spot-level resolution. The mIF platform PhenoCycler (from Akoya Biosciences)^18^ enables protein profiling (throughput ∼ 40) at a 0.6 *µm* cellular resolution. Despite these differences, the underlying data structure remains largely consistent across technologies and platforms, comprising a collection of spatial locations (from single or multiple samples) with observed expression or intensity of various molecules. Consequently, common biostatistical questions arise, centering the spatial dynamics of molecules within the complex tissue or tumor microenvironment (TME)^19–23^.

In the context of ST datasets, identifying spatially variable genes (SVGs), i.e., the genes exhibiting spatially structured expression patterns across the tissue, has gained significant attention^24–38^. It enables critical downstream analyses such as discovering potential biomarkers and defining tissue regions that influence cellular differentiation and function^39–42^. Analogously, for mIF or imaging mass cytometry (IMC) datasets, innovative methods^43–51^ have been proposed to understand the spatial distribution of immune cell types (defined by binarizing the expression profile of immune proteins) across the TME. Building upon this univariate framework, which typically analyzes one molecule at a time, another widely investigated problem has been spatial domain detection, i.e., deconvolving the tissue into distinct, spatially contiguous neighborhoods based on multivariate gene expression (ST)^52–63^ or immune cell type composition (mIF)^64–69^. It aids mapping the molecular and functional landscape of tissues, elucidating disease progression, and guiding targeted therapies^70–72^. While some of the referenced methods can be adapted for use with MSI datasets, it is important to underscore the lack of sophisticated spatial functionalities of the existing bioinformatics toolboxes^73–76^.

While univariate and multivariate spatial analyses have garnered significant attention, a critical intermediate task remains underexplored: bivariate spatial co-expression analysis of molecular pairs at both “local” (spot/cell-specific) and “global” (tissue-wide) levels, aimed at precisely characterizing the spatial interaction or binding pattern of any two molecules throughout the tissue plane. To emphasize the importance of such an analysis, we review the concepts of cell-cell communication (CCC)^77–80^. CCC is a fundamental biological process through which cells exchange information via direct contact or signaling molecules (ligands) binding to receptor molecules present on the same or different cells. It regulates essential biological functions, including tissue development^81^ and immune responses^82^, and its disruption has been implicated in the onset and progression of cancer^83^. Autocrine, juxtacrine, and paracrine signaling are three major pathways of CCC^84^. In autocrine signaling, ligands released by a cell bind to receptors on the same cell, while in juxtacrine and paracrine signaling, the ligands target adjacent and nearby cells. The study of ligand-receptor interactions (LRI), which involves identifying gene pairs (ligands and receptors) that show coordinated upregulation or downregulation across groups of cells, has become a fundamental approach for inferring CCC from single-cell RNA sequencing (scRNA-seq) datasets^85–93^. However, these approaches are prone to false positive interactions due to the lack of spatial context in scRNA-seq datasets, treating distant cell pairs similarly to nearby ones^94–96^, which potentially leads to an overestimation of juxtacrine and paracrine signaling. ST datasets offer a natural avenue for improvement by enabling spatially constrained LRI analysis.

A limited number of tools exist for spatial LRI analysis or, more broadly, for assessing bivariate spatial co-expression of molecules in ST or MSI datasets. It should be emphasized that bivariate co-expression can manifest in two ways: (a) joint over- or under-expression within the same cells (correlation) and (b) joint over- or under-expression in neighboring cells (cross-correlation^97^). Some relevant methods include MERINGUE^98^, Giotto^99^, SpaGene^100^, SpaTalk^101^, SpatialDM^102^, CellChat V2^103^, LIANA+^104^, and Copulacci^105^. We skip the approaches that jointly analyze multiple LR pairs^106,107^. Methods such as MERINGUE, Giotto, and SpaTalk provide only a global summary of spatial co-expression across a tissue, whereas others also offer local (spot/cell-specific) estimates. Let the standardized expression of two genes (*m, m*^′^) be *X*^*m*^(*s*) and *X*^*m*′^ (*s*) at location *s* for *s* ∈ {*s*_1_, …, *s*_*n*_}, and *X*^*m*^ = (*X*^*m*^(*s*_1_), …, *X*^*m*^(*s*_*n*_))^*T*^, *X*^*m*′^ = (*X*^*m*′^ (*s*_1_), …, *X*^*m*′^ (*s*_*n*_))^*T*^ . For a global summary of spatial co-expression, MERINGUE and SpatialDM leverage a popular geospatial metric termed the bivariate Moran’s *I* (*I*_*BV*_)^108,109^, interpreted as the Pearson correlation between one variable and the spatial lagged version of the other^110–112^. Mathematically, *I*_*BV*_ ∝ (*X*^*m*^)^*T*^ *WX*^*m*′^, where 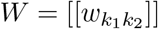 is the spatial weight matrix that controls the spatial lagging. As *W*, MERINGUE uses a binary adjacency matrix based on the Delaunay triangulation^113^ of the spatial locations (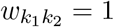 if locations 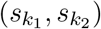 are connected, or 0 otherwise). SpatialDM uses a kernel covariance matrix or Gram matrix^114^ based on the *L*_2_ distance between locations 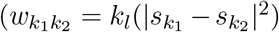, where *k*_*l*_ is a kernel function with lengthscale parameter *l*^115^). For local estimates of spatial correlation, SpatialDM considers the bivariate local Moran’s 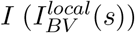 based on the local indicators of spatial association (LISA) approach^116^. The LIANA+ toolbox implements SpatialDM and introduces a similar spatially weighted cosine similarity index. Of note, a newer package named Voyager^117^ considers Lee’s *L* statistic^110^, which has a slightly different formulation than *I*_*BV*_ . A critical yet often overlooked aspect of ST data analysis is that gene expression, measured in terms of unique molecular identifier (UMI) count, is inherently a discrete random variable (RV). However, the above methods assume normality upon a variance-stabilizing transformation^118–120^, which may obscure true signals and have been widely criticized both within the ST literature^26,55,121,122^ and in broader contexts^123–126^. Addressing this issue, Copulacci models a pair of genes as bivariate Poisson-distributed RVs, with their correlation in spatially adjacent cells captured using a Gaussian copula^127^. For inference, these methods typically rely on a permutation test^128^.

Bivariate Moran’s *I* (*I*_*BV*_) and Lee’s *L* statistic, as implemented in MERINGUE, SpatialDM, LIANA+, and Voyager, are primarily recommended as exploratory metrics for assessing cross-correlation rather than as rigorous hypothesis testing tools^129,97^, in traditional spatial statistical literature. In simulation studies (see Section 2.2), we have shown that even when two variables are independently simulated with certain spatial covariance structures, the unmodeled spatial autocorrelation introduces a confounding effect on the bivariate association, leading to significantly inflated Type I error rates. A similar issue is well documented, as extensive literature highlights the limitations of using simple Pearson correlation to assess dependencies between two variables in the presence of spatial autocorrelation^130–134^. By extension, since *I*_*BV*_ and Lee’s *L* are both fundamentally based on Pearson correlation between spatially lagged variables, they may be susceptible to similar pitfalls. Furthermore, these spatially weighted association indices, being model-free, are unable to seamlessly adjust for cell-level covariates such as cell type, a limitation also present in Copulacci. As a side note, mapping to the aforementioned CCC pathways, Pearson correlation between ligand and receptor can be interpreted as a proxy for autocrine signaling, while cross-correlation may reflect a combination of juxtacrine and paracrine signaling.

We approach the bivariate spatial co-expression detection as a generalized linear regression problem, modeling a molecule *m* as the outcome and the other molecule *m*^′^ as the predictor (see Section 4). For ST datasets, gene expression or UMI count is modeled as an overdispersed negative binomial (NB)-distributed RV^135^, while an alternative Gaussian model is considered for continuous cases. The regression coefficients, both intercept 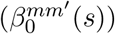 and slope 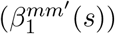, are assumed to vary across locations (*s*) exhibiting spatial dependency. Known as the spatially varying coefficients (SVC) model^136^, this framework provides exceptional flexibility and precision in capturing locally changing co-expression patterns through 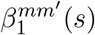. A large positive 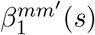 suggests strong positive co-expression at location *s*, i.e., joint up or down-regulation, whereas a large negative value indicates avoidance or repulsion. The average of 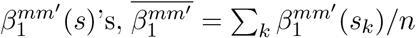 provides a summary of the global co-expression pattern. Similar models have been widely used in fields such as disease mapping^137,138^, econometrics^139,140^, ecological studies^141,142^, and neuroimaging research^143,144^. In the Bayesian paradigm, the spatial dependency between 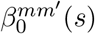 ‘s and 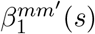 ‘s is typically modeled using a conditional autoregressive (CAR)^145–147^ or Gaussian process (GP) priors^148,97,149^. In contrast, we introduce a locally adaptive spatial Gaussian Markov random field (GMRF) prior^150^ based on the concepts of fusion penalties^151–153^ and horseshoe prior^154–156^, extending a related work in the frequentist setup^157^. Briefly, the prior incorporates the spatial similarity between two adjacent locations, 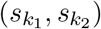, by encouraging 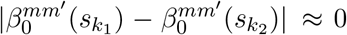 and 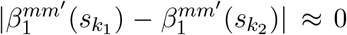. We parameterize spatial adjacency primarily with the minimum spanning tree (MST)^158–160^, following Li et al. (2019), which provides cycle-free, globally economical connectivity by minimizing total edge weight. In practice, we find that modestly denser graphs, e.g., *k*-nearest neighbors with small *k*, can yield improved performance. We evaluate the proposed method SpaceBF against established approaches under realistic simulation scenarios, demonstrating high specificity and power. For broader applicability, we also benchmark our prior against the standard intrinsic CAR (ICAR) and a stochastic partial differential equation (SPDE)-based Matérn prior^161,162^, where SpaceBF consistently outperforms both alternatives. SpaceBF is applied to three real datasets: a) an ST dataset on cutaneous melanoma^39^ for spatial LRI analysis, b) an ST dataset on cutaneous squamous cell carcinoma^163^ for keratin-interaction analysis, and c) a spatial proteomics dataset on ductal carcinoma in situ (DCIS) from the Medical University of South Carolina (MUSC) for peptide co-localization analysis. An efficient R package with different choices of priors and models is available at https://github.com/sealx017/SpaceBF/.

## 2 Result

### 2.1 Real data analysis

We use the MST as the spatial adjacency graph for the real datasets in the main text and provide complementary *k*-NN–based results in the Supplementary Material.

#### 2.1.1 Melanoma ST dataset

We analyzed a cutaneous melanoma dataset^39^ from a long-term survivor (10+ years), collected using the ST technology^4^, comprising 293 spots, each 100 *µm* in size and at a 200 *µm* center-to-center distance. There are 16,148 genes, forming 1,180 known ligand-receptor (LR) pairs as available from CellChatDB^88^. There are three major pathologist-annotated regions as seen in the histology image (Fig. 1A), collected from Thrane et al. (2018), and 6 major cell types (Fig. 1B) predicted using the RCTD^164^ package based on overall gene expression^102^. After filtering out genes with extremely low expression (*<* 0.2 × 293 ≈ 59 reads), 161 LR pairs remain, which were examined using our method SpaceBF, without adjusting for any covariates. To briefly summarize the SpaceBF workflow, it first constructs an MST based on the spatial coordinates of the spots (Fig. 1C). Then, for every LR pair: (*m*^′^, *m*), it considers Eq. 2 with the receptor expression as *X*^*m*^(*s*_*k*_) and the ligand expression as *X*^*m*′^ (*s*_*k*_), and *s*_*k*_ representing a spot. Following parameter estimation via a Markov Chain Monte Carlo (MCMC) procedure, the framework performs two hypothesis tests to assess the significance of spatial co-expression at both global and local levels (see Section 4.3). Using the global test in this dataset, SpaceBF identified 53 LR pairs at a significance level 0.05 (33 at an FDR of 0.1). The estimated slope surface 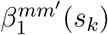 of different LR pairs exhibits distinct patterns. To highlight these differences, we classify the detected LR pairs into 3 major patterns (Fig. 1E) based on hierarchical clustering^165^ of the standardized vector 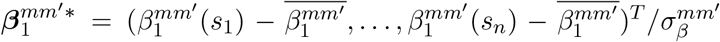, where 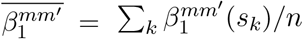 and 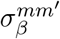 are the tissue-wide average and the SD of estimated 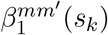 ‘s, respectively. 20 LR pairs follow pattern 1, while 22 and 11 LR pairs correspond to patterns 2 and 3, respectively. Similarly, the spots are grouped into 4 clusters based on the spot-level vectors of slopes corresponding to the 53 detected LR pairs (Fig. 1D). It is evident that clusters 1 and 3 correspond to the melanoma region, while clusters 2 and 4 loosely correspond to the stroma and lymphoid regions, respectively. Returning to the LR patterns, in Fig. 1F, the LR pairs are arranged sequentially from pattern 1 to 3, highlighting the enrichment of their interaction in three major cell types. For example, 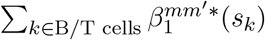 represents the enrichment within B/T cells relative to the average enrichment 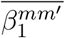 and scaled by the SD. The levels “highest,” “medium,” and “lowest” indicate the degree of enrichment, with “highest” corresponding to the greatest or most positive enrichment and so on. The majority of LR pairs following pattern 1 exhibit higher or more positive interaction in B/T cells within the lymphoid region (some in CAF cells) and more negative interaction (avoidance or repulsion) in the melanoma region or cells. Pattern 2 mostly corresponds to LR pairs with the highest enrichment in CAF cells, while pattern 3 clearly corresponds to the pairs with the highest enrichment in melanoma cells. Next, we investigate the biological relevance of the estimated slope surfaces for a selected set of LR pairs. The LR pair (IGF2, IGF1R)^166^ corresponds to pattern 1 and demonstrates a negative association overall, with an estimated average slope of 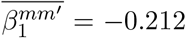, and the *p*-value = 0.024, which is consistent with a visual inspection (Fig. 1G). It could indicate a lack of binding between these genes, which would be a generally favorable factor for the survivor^167^. Setting the insignificant 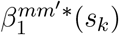 values to 0 based on the local test, the negative interaction found in the melanoma region has the highest credibility. The pair (PTPRC, CD22)^168^ follows pattern 2, with 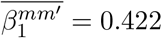 and *p*-value of 6.28 × 10^−6^. PTPRC, also known as CD45, is a facilitator of T-cell receptor (TCR) and B-cell receptor (BCR) signaling^169^, while CD22 is primarily an inhibitor of BCR signaling^170^. Their overall positive co-expression, particularly in the lymphoid region, is likely associated with a balanced B cell regulation, helping to prevent autoimmunity and promoting lymphoid growth in other regions as part of the immune response. The final LR pair we discuss is (SPP1, CD44)^171^, which follows pattern 3, exhibiting a highly positive overall co-expression with 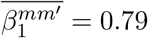 and *p*-value of 1.03 × 10^−6^. This strong interaction displays a decreasing gradient from the melanoma region to the lymphoid region, which aligns with its known role in dysregulated cytoskeletal remodeling^172^, facilitating melanoma cell invasion into surrounding tissues.

**Figure 1:**
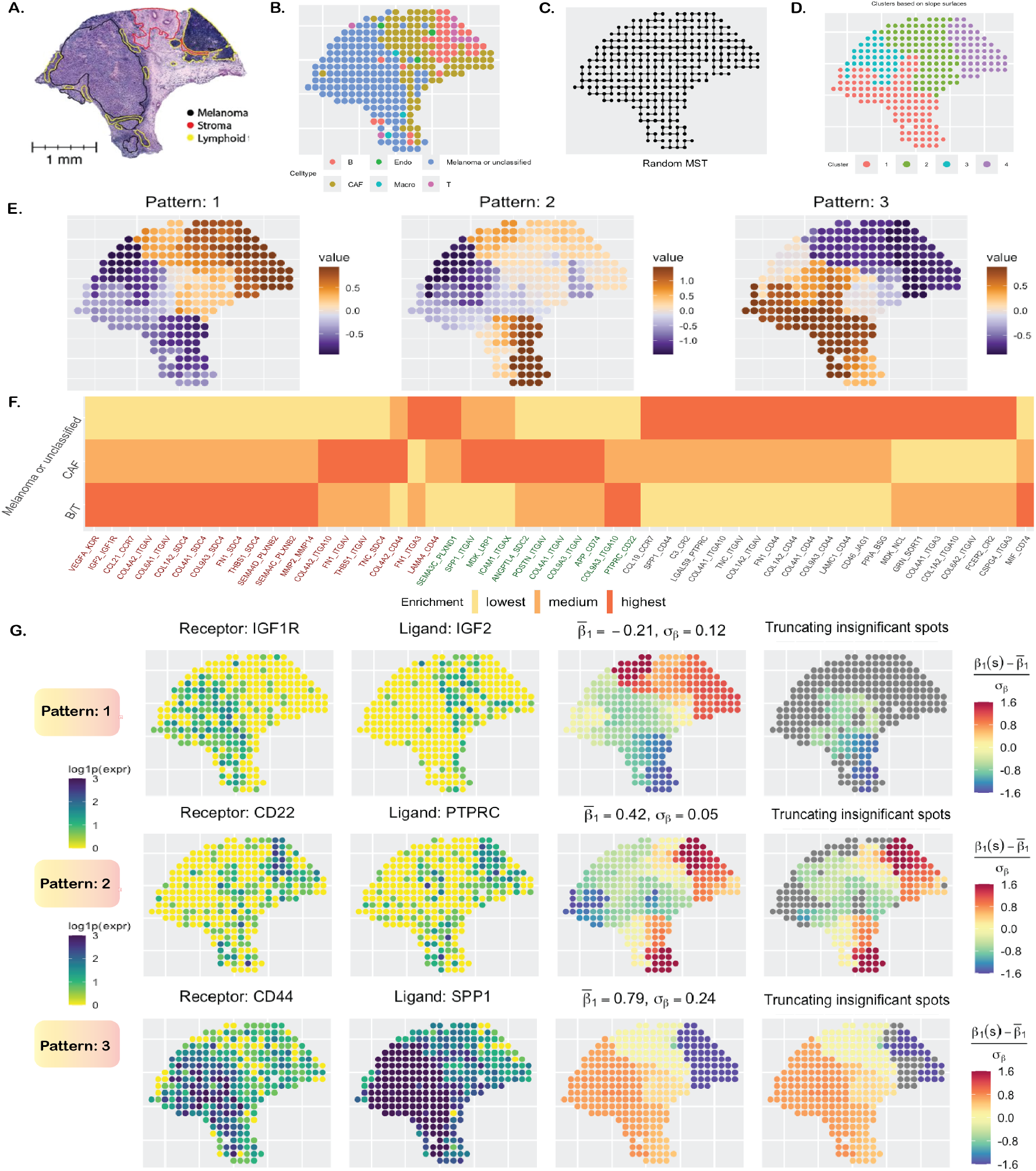
Cutaneous melanoma data analysis. **A**. Annotated H&E-stained image. **B**. Cell types based on gene expression. **C**. Minimum spanning tree (MST) capturing the spatial structure. **D**. Clustering of spots based on centered and scaled estimates of slope surfaces of 53 statistically significant LR pairs. **E**. The three main spatial patterns of the estimated surfaces. **F**. Enrichment of LR interactions in three major cell types, with LR names arranged and color-coded according to their respective patterns. **G**. The first two columns show the expression of three LR pairs. The third column displays the centered and scaled slope surfaces. In the fourth column, insignificant spot-level slope estimates are greyed.

#### 2.1.2 cSCC ST dataset

We analyzed a cutaneous squamous cell carcinoma (cSCC) dataset^163^ on a patient sample with a histopathologic subtype of “moderately differentiated” cSCC^173^. The dataset was collected using the ST technology with 621 spots, each of size 110 *µm* and a center-to-center distance of 150 *µm*. There are 16, 643 genes of which 45 are keratins (14 after filtering low-count genes, *<* 0.2 × 621 ≈ 124 reads). These keratins can be classified into two types: 1) Type I, which includes KRT10, KRT14–KRT17, and KRT23, and 2) type 2, which includes KRT1, KRT2, KRT5, KRT6A, KRT6B, KRT6C, KRT78, and KRT80. The keratins pair together to form intermediate filaments, providing structural support to epithelial cells^174^. In the context of cSCC and other carcinomas, keratins are emerging as highly significant targets for therapeutic intervention^175–177^. Of note, some of the keratins belong to the GO term: “keratinocyte differentiation” (GO:0030216) and were reported to exhibit strong spatial correlation in an earlier work^178^ involving the same dataset. We utilized SpaceBF to investigate the binding between Type I and type 2 keratins, resulting in a set of 48 keratin pairs. In the histology image (Fig. 2A), the deep blue areas at the top and left sides correspond to tumor regions, while the whitish region at the bottom represents a non-tumor region possibly composed of keratinized layers and stroma^179^. However, the tumor and non-tumor regions are not clearly delineated, a feature characteristic of moderately differentiated cSCC, though the spatial clusters obtained using the BayesSpace package^52^ on the transcriptome-wide gene expression profile (Fig. 2C) partially elucidate this distinction. The constructed MST is shown in Fig. 2B. Using the global test, SpaceBF identified 39 keratin pairs at a significance level of 0.05 (41 at an FDR *<* 0.1), suggesting that most pairs bind to each other, albeit to varying degrees. Similar to the earlier analysis, we classify the detected slope surfaces into 3 major patterns (Fig. 2D) based on hierarchical clustering of the standardized vector 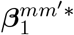. We represent the keratin pairs as bipartite graphs between Type I and 2 keratins under each pattern (Fig. 2E). One important observation is that type 2 keratins KRT6A, KRT6B, and KRT6C are isoforms of keratin 6^180^ and thus, co-express highly, meaning their binding patterns with any specific Type I keratin should be similar, as correctly identified by SpaceBF. For instance, the slope surfaces of KRT10 with KRT6A, 6B, and 6C all align with pattern 1, while the slope surfaces of KRT16 with KRT6A, 6B, and 6C all correspond to pattern 2. This consistency underscores the reliability of SpaceBF in identifying true local patterns. In Fig. 2F, we present the estimated slopes for KRT17, which is a well-established therapeutic target in various cancers^181–183^, binding with three type 2 keratins: KRT80 (pattern 1, *p*-value = 0.006), KRT78 (pattern 2, *p*-value = 0.03), and KRT6B (pattern 3, *p*-value = 5.59 × 10^−6^). Notably, the average slope estimates for KRT17-KRT80 and KRT17-KRT78 interactions are small (≈ 0.1), whereas for KRT17-KRT6B, the average slope is substantially higher at 0.82, with the highest local estimates observed mostly in tumor regions. These trends are also evident from the individual expression profiles provided in Fig. 2F. Although the expression patterns of KRT80 and KRT78 appear similar, a closer examination reveals that KRT80 exhibits a thicker band of expression on the left, specifically within the tumor regions. This distinction contributes to the difference in co-expression patterns of KRT17-KRT80 and KRT17-KRT78. As previously noted, both association levels are low, also indicated by the small number of significant spots identified by the local test, 72 and 49, respectively.

**Figure 2:**
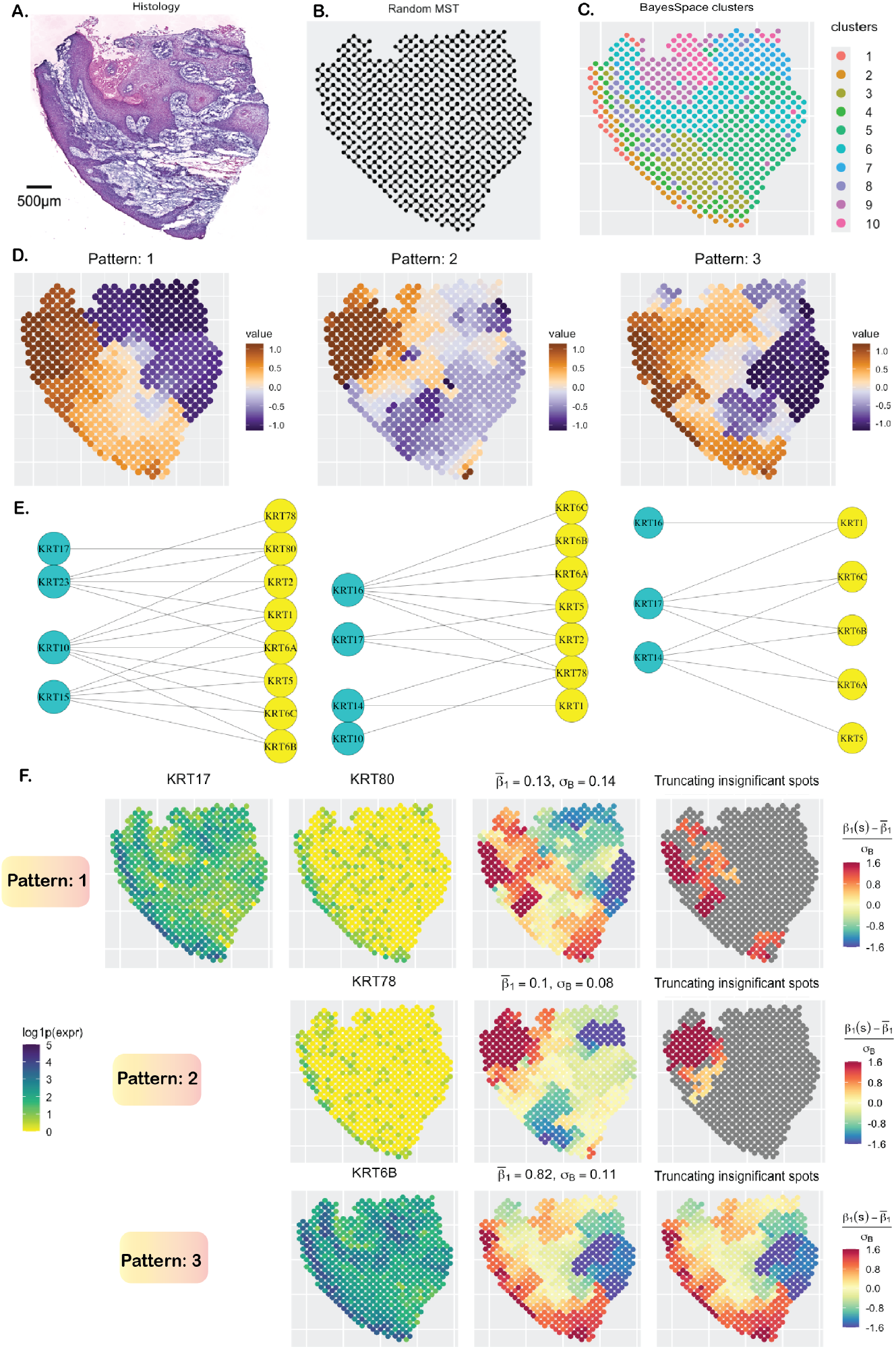
Cutaneous squamous cell carcinoma data analysis. **A**. H&E-stained image. **B**. MST capturing the spatial structure. **C**. Spatial clusters obtained using the BayesSpace package. **D**. The three main spatial patterns of the estimated surfaces. **E**. Bipartite graphs between Type I and type 2 keratins based on their spatial pattern. **F**. Study of the binding between the Type I keratin KRT17 and three different type 2 keratins, with each slope surface exhibiting a unique spatial pattern. The insignificant spot-level slope estimates are greyed in the last column.

#### 2.1.3 DCIS proteomics dataset

We analyzed a single-sample ductal carcinoma in situ (DCIS) dataset collected using the MALDI MSI spatial proteomics platform, as part of an ongoing study at the MUSC aimed at defining the proteomic landscape of DCIS and invasive breast cancer (IBC), in terms of collagen peptides and immune cell types. DCIS is marked by the abnormal growth of malignant epithelial cells confined to the breast’s milk ducts, without invading the surrounding stromal tissue. While prognosis is excellent, around 20−40% of diagnosed DCIS progress to IBC^184,185^. Understanding proteomic co-localization within the extracellular matrix (ECM) of a DCIS tissue is crucial for assessing progression risk and predicting therapeutic response, as the ECM plays a key role in regulating tumor cell proliferation, migration, and survival^186^. In this dataset, there are 5,548 tissue spots and 12 ECM peptides, whose 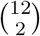 pairwise interactions were of our interest. As the peptide expression is continuous-valued, we used the Gaussian model of SpaceBF for this analysis (Eq. 1). Seven of the peptides are derived from the COL1A1 gene, while the remaining peptides originate from COL1A2, COL3A1, and FN1 (Fig. 3C). From the histology image (Fig. 3A), the stromal ECM can be identified by the light pink staining of fibrous connective tissue, while epithelium regions are highlighted in deep blue. Using hierarchical clustering, we group the standardized slope surfaces 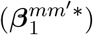 into three patterns (Fig. 3B), and the spots into three clusters based on the spot-level vectors of standardized slopes (Fig. 3D). While the differences among the three patterns are subtle, the spot clusters are well-defined and spatially distinct: the red cluster (cluster 3) aligns with stromal regions, the green cluster (cluster 2) with epithelial regions, and the light blue cluster (cluster 1) is a mixture of both. It is important to note that the MSI image has substantially lower resolution compared to the histology image, making one-to-one correspondence between the two inherently challenging. Patterns 1 and 2 (Fig. 3B), which visually resemble each other, both suggest strong co-localization of the associated peptide pairs in the stroma. This is expected, as all of these peptides are known to constitute the stromal ECM. The tree diagram in Fig. 3C shows the hierarchical relationships between the peptide pairs, with their patterns indicated on the right. The module highlighted by the yellow box includes pairs involving peptide 1125 (from COL1A2) and 7 other peptides. From Figs. 3E and 3F (top row), peptides 1125, 1212, 1386, and 1681 (from the module) show pronounced co-expression in the stromal region. Correspondingly, the estimated slope surfaces (Fig. 3F, bottom row) for the pairs (1125, 1212), (1125, 1386), and (1125, 1681) all fall under pattern 1, but the association strength is notably higher for (1125, 1212):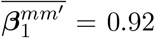, compared to 0.72 and 0.68 for the other two. Although the existing literature on these interactions is limited, the findings will inform future comparative analyses of ECM compositions across DCIS subtypes and stages of progression^187,188^.

**Figure 3:**
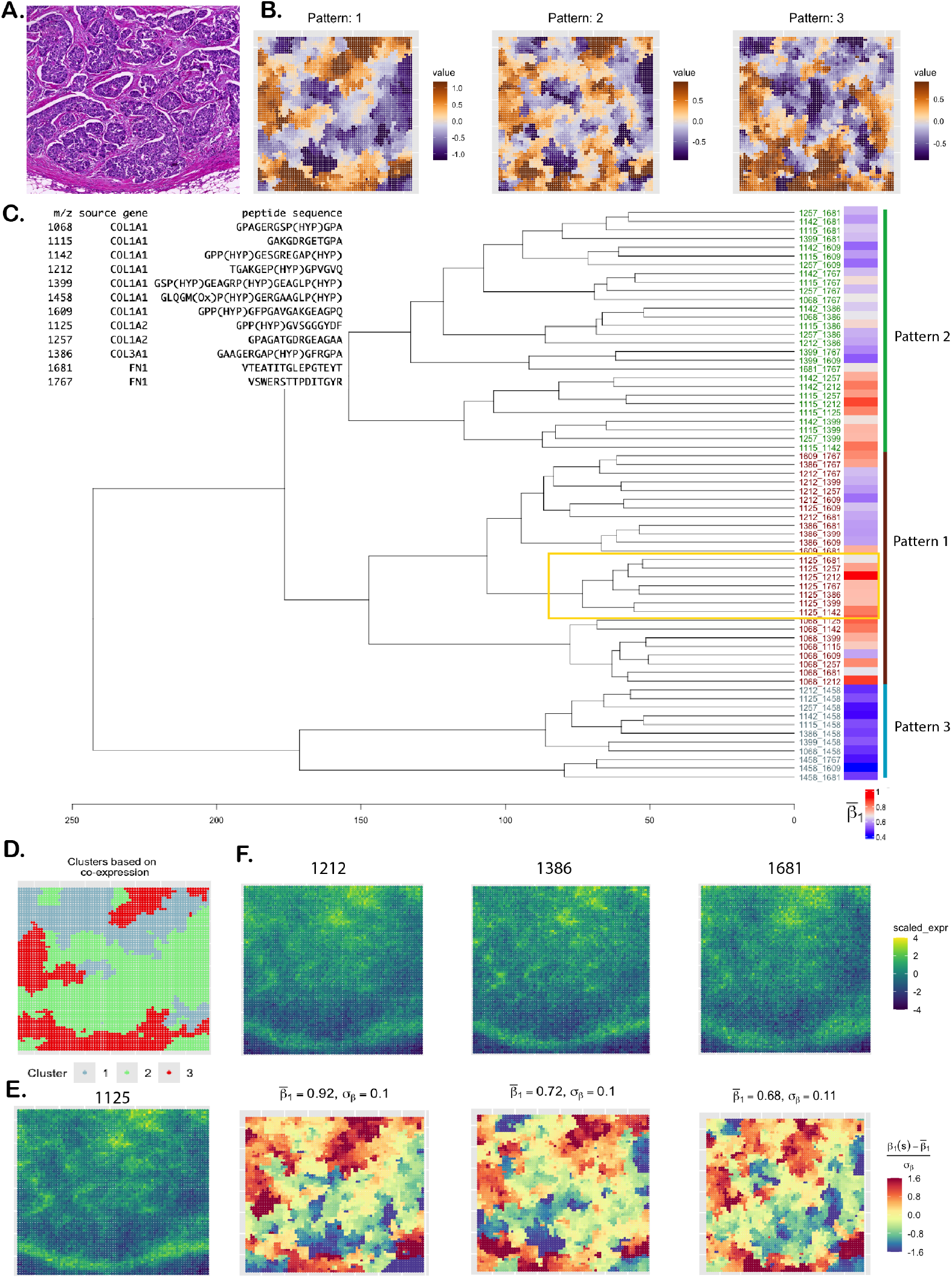
DCIS data analysis. **A**. H&E-stained image. **B**. Patterns of standardized co-expression (slope) of 66 peptide pairs (from 12 peptides). **C**. Peptide description and dendrogram corresponding to the patterns. Mean slope estimates are presented as a heatmap on the right. **D**. Clustering of spots based on the slope surfaces. **E**. Scaled expression of the peptide 1125 forming the yellow-bordered module in the dendrogram. **F**. Scaled expression of three peptides belonging to the same module (top row) and their spatial co-expression with peptide 1125 (bottom row).

### 2.2 Simulation studies

#### 2.2.1 Simulation design 1: comparison between global methods under linear relation

We consider the spatial coordinates (*n* = 293) from the cutaneous melanoma dataset. In simulation design 1, one NB-distributed random variable (RV), **X**^*m*′^ is generated using a Gaussian copula with a spatial covariance matrix *H* based on an exponential kernel (for varying lengthscale *l*) and the *L*_2_ distance. Another NB-distributed RV, **X**^*m*^ is then generated using the NB model from Eq. 2 with a constant slope 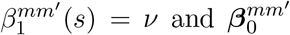 simulated using a Gaussian process (GP) model^97^ with the spatial covariance matrix *H*. More details on the design are provided in Section 4.4.1. From Fig. 4A, we notice how the structure of *H* changes as the lengthscale *l* varies. The off-diagonal elements of *H* 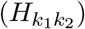 can range between 0 and 1. When *l* = 0.6, only the nearest locations (*k*_1_, *k*_2_) exhibit high 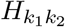 with most of the other values being close to 0. In contrast, for *l* = 18, the majority of location pairs have high 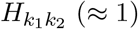, inducing an exceptionally strong spatial autocorrelation in both variables. To visibly understand how *ν* might affect the relationship between **X**^*m*^ and **X**^*m*′^, in Fig. 4B, we show the spatial expression of **X**^*m*^ for the same **X**^*m*′^ but three different values of *ν*, {−0.75, 0, 0.75}. It is somewhat evident that nonzero *ν*’s result in a visibly positive or negative association, while *ν* = 0 produces a random pattern of **X**^*m*^. In Fig. 4C, we show the Type I error (*ν* = 0) and power (*ν* ≠ 0) comparison of the different methods, including SpaceBF, for three values of the lengthscale *l*. When *l* = 3.6, both variables exhibit considerable spatial autocorrelation, yet SpaceBF maintains the correct Type I error. In contrast, all other methods suffer from inflated Type I errors. Notably, simple Pearson correlation, while still inflated, performs better in controlling Type I error compared to methods based on bivariate Moran’s *I* or Lee’s *L*. Although SpaGene does not rely on these traditional metrics, it still fails to control Type I error. The issue becomes more pronounced as *l* increases. Notably, Lee’s *L* exhibits the highest inflation in the majority of cases. SpaceBF also retains a high detection power throughout all three cases. Although the power declines slightly for the largest *l*, as expected, due to a decrease in effective sample size from increased spatial autocorrelation. In summary, the simulation effectively demonstrates the specificity and power of our method.

**Figure 4:**
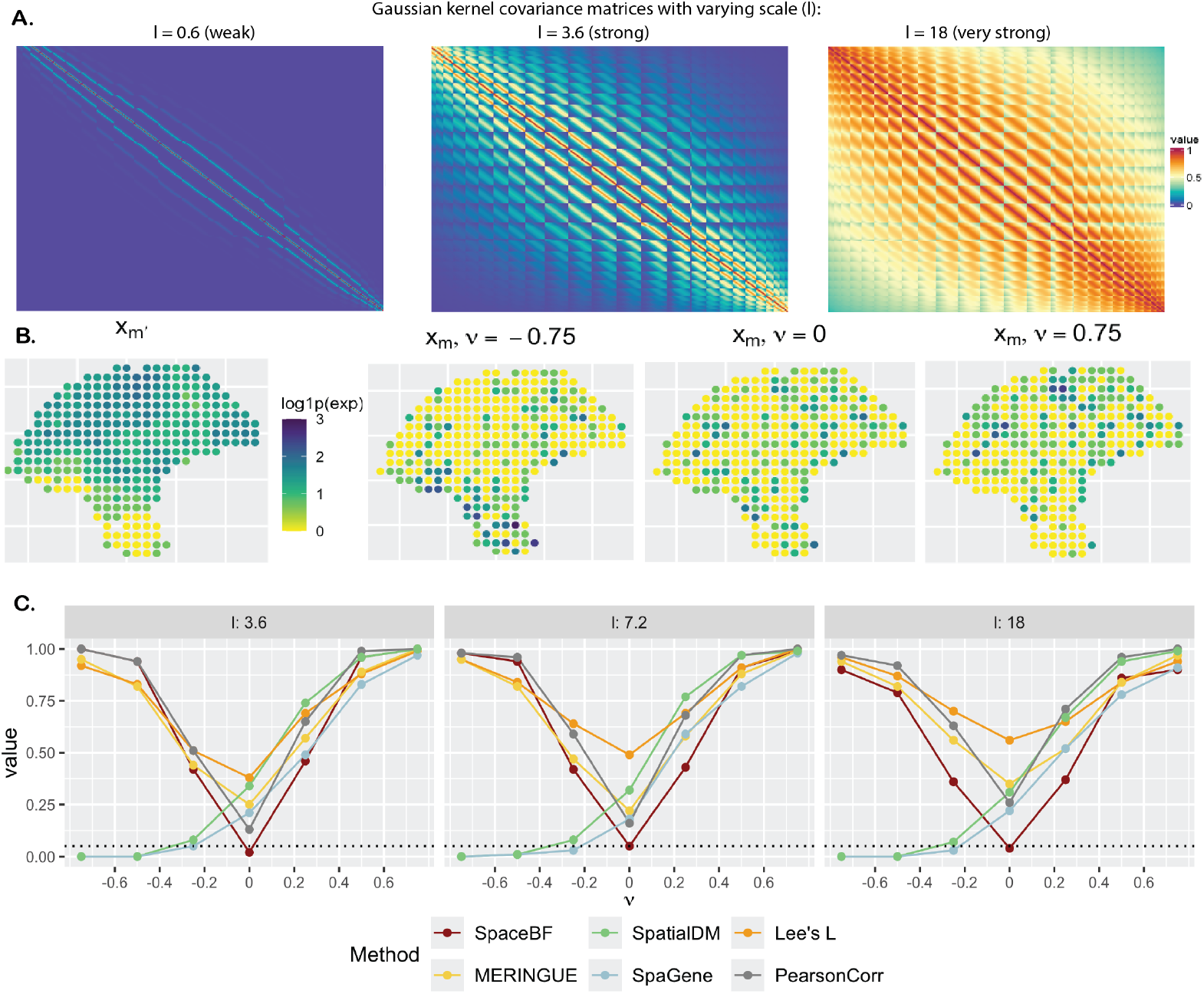
Comparison of global tests under the simulation design 1. **A**. Heatmap of the spatial covariance matrix *H* with an exponential kernel and *L*_2_ distance, for varying values of the lengthscale parameter *l*. **B**. Simulated **X**^*m*^ based on Eq. 2, for a fixed **X**^*m*′^ but different values of the constant slope 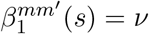. **C**. Performance of the methods in terms of Type I error (*ν* = 0) and power (*ν* ≠ 0). The dotted line represents the significance level 0.05.

#### 2.2.2 Simulation design 2: comparison between global methods under non-linear relation

In simulation design 2, we generate (**X**^*m*^, **X**^*m*′^) jointly as bivariate spatially correlated NB-distributed RVs. This setup is more complex than the previous one, as the association is non-linear and driven by the Kronecker product-based spatial covariance structure (see Section 4.4.2). The methods, except SpaceBF, perform poorly in terms of the Type I error for *l* ≥ 1.8. When the spatial autocorrelation is the weakest (*l* = 0.6), Pearson correlation performs well as expected, while spatially weighted indices still show a slight inflation. SpaceBF achieves controlled Type I error and steady detection power across varying *l*’s. As earlier, the power decreases as the effective sample size decreases. Together, these two simulation designs demonstrate SpaceBF’s robustness under complex data generation processes. Finally, we argue that incorporating 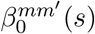 in SpaceBF accounts for the spatial autocorrelation of variable *m*, thereby mitigating bias in the association analysis of (*m, m*^′^). As previously noted, Pearson correlation is already recognized to be suboptimal in such scenarios, and spatially weighted indices, essentially Pearson correlation between spatially lagged variables, thus also remain susceptible to spurious detections. Recall that MERINGUE uses a binary spatial weight matrix *W* based on the Delaunay triangulation, while SpatialDM uses a continuous spatial weight matrix having a similar form as *H* for a particular choice of the lengthscale. Lee’s *L* is based on a binary *k*−NN network in our study. Hence, the performance of these methods could be sensitive to choices of the spatial weight matrices, i.e., different networks or lengthscale *l* values. SpatialDM and SpaGene focus solely on the joint over-expression of molecules, neglecting joint under-expression. As a result, for most values of *ν <* 0, these methods show almost no detection power.

#### 2.2.3 Simulation design 3: comparison between spatial priors under SVC framework

While the spatial horseshoe (HS) prior is introduced on a minimum spanning tree (MST), it can be placed on any spatial backbone (e.g., Delaunay or *k*-NN graphs), albeit with a potential risk of oversmoothing. This simulation study evaluates how graph choice affects HS performance. A Delaunay network is substantially denser than an MST, whereas a *k*-NN network can serve as a middle ground for small *k*. In Fig. 6, HS-MST denotes HS on the MST (the original SpaceBF setting used in previous simulations and applications), HS-Del denotes HS on the Delaunay graph, and HS-*k*NN denotes HS on a *k*-NN graph with *k* = 3. As noted in the Methods section, the ICAR prior is a special case of the HS prior; we therefore include ICAR-Del and ICAR-*k*NN for comparison. For completeness, we also consider a stochastic partial differential equation (SPDE)^161^-based NB SVC model implemented in the efficient `R` package `sdmTMB`^162^, which uses a Matérn prior: sdmTMB-Matérn1 uses a denser mesh (cutoff = 1), and sdmTMB-Matérn2 uses a coarser mesh (cutoff = 1.5), see the Supplementary Material for a visual comparison.

**Figure 5:**
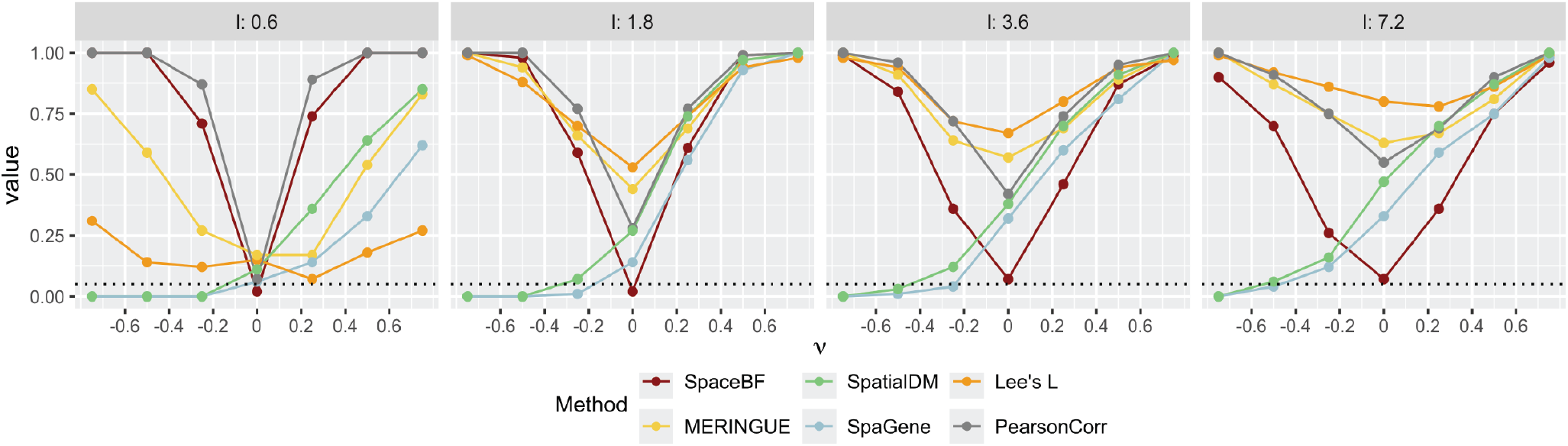
Comparison of global tests under simulation design 2 for lengthscale *l* between {0.6, 1.8, 3.6, 7.2}.

**Figure 6:**
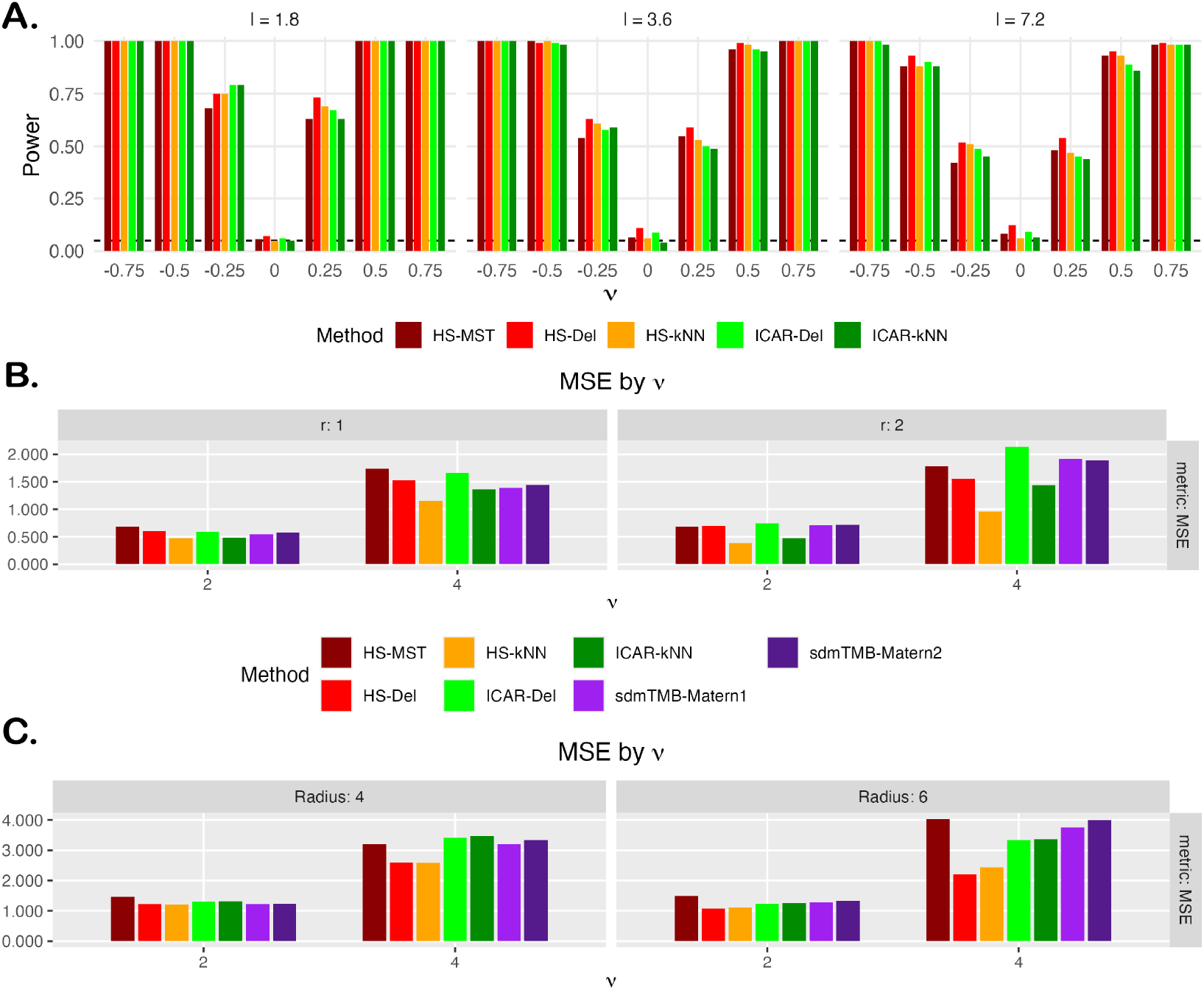
**A**. Power comparison of spatial priors under simulation design 2 for lengthscale *l* between {1.8, 3.6, 7.2}. **B**. MSE comparison of spatial priors under simulation design 3, linear partition boundary. **C**. MSE comparison of spatial priors under simulation design 3, circular boundary. In panel A, sdmTMB models are omitted due to recurrent convergence issues.

We first revisit the simulation design 2 to assess global performance under a nonlinear model. As shown in Fig. 6A, all priors perform similarly across lengthscales, with a modest inflation in Type I error for HS-Del and ICAR-Del. This aligns with the oversmoothing tendency of denser backbones, which can spuriously propagate a small local positive association (e.g., confined to one corner of the tissue) across the entire domain. The overall similarity among priors is unsurprising in this particular simulation setting: with co-expression effectively constant over space, the additional flexibility of HS (an adaptive precision matrix) is not meaningfully exercised. Notably, `sdmTMB`-based methods failed to converge in several scenarios (and are therefore omitted here), especially when the outcome contained a high proportion of zeros. We attribute these failures to the near-vanishing curvature of the NB log-likelihood under heavy zero counts, which potentially destabilizes the Laplace-based frequentist optimization; in such cases, a Bayesian approach via `R-INLA`^189^ may be preferable.

Next, we assess each prior’s ability to recover *spatially varying* slopes under the linear and circular boundary designs in Section 4.4.3. In Fig. 6B, HS-*k*NN achieves the lowest mean squared error (MSE) across both linear-boundary settings (both values of *r*) and for both effect sizes (*ν* = 2 and *ν* = 4). The gain is most pronounced at *ν* = 4, where its MSE is roughly half of the competing methods. By contrast, HS-MST, sdmTMB-Matérn1, and sdmTMB-Matérn2 generally perform the worst. For the circular boundary (Fig. 7C), HS-Del and HS-*k*NN perform similarly and outperform the remaining approaches. Overall, the MSE results favor the HS prior over standard ICAR models on generic spatial graphs and over the Matérn (SPDE-based) alternative. Because HS-MST is consistently outperformed by HS-*k*NN, we recommend using a denser graph such as *k*NN in typical applications. Moreover, Figs. 6C and 7C show that HS-*k*NN best preserves sharp slope changes at boundaries, whereas ICAR methods (especially ICAR-Del) and sdmTMB methods (notably sdmTMB-Matérn2 with a coarser mesh) tend to blur these transitions, highlighting the HS prior’s ability to capture subtle co-expression shifts in the TME. Finally, while HS-Del can occasionally perform well, it may oversmooth and produce spurious detections, cautioning against the use of overly dense graphs such as Delaunay.

**Figure 7:**
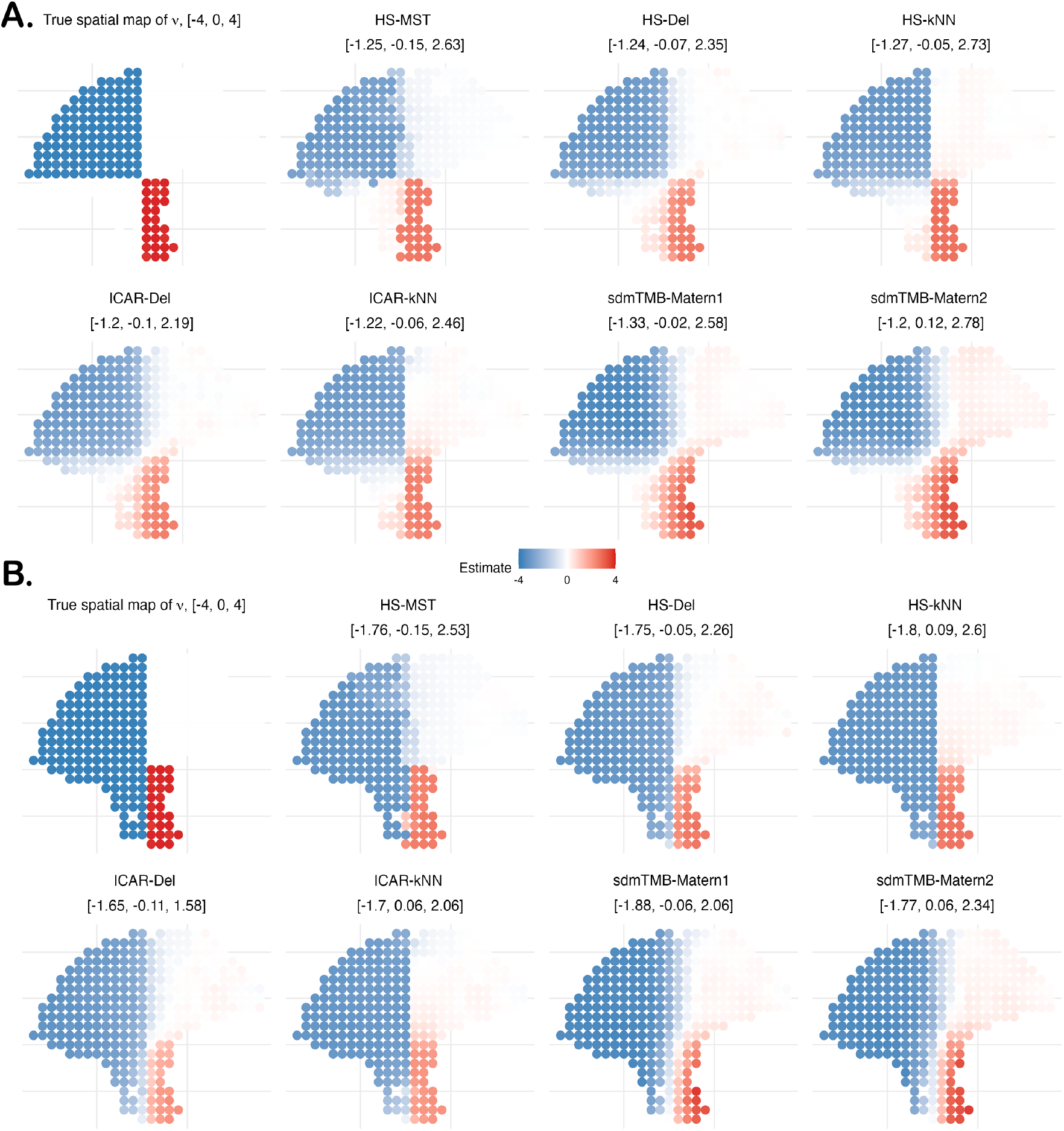
SVC simulation based on linear partition boundary from Section 4.4.3 with effect size *ν* = 4. The boundary pattern changes based on the parameter *r*: **A**. *r* = 1, more zeros on the lower left, and **B**. *r* = 2, more negative slope values on the lower left.

## 3 Discussion

We have developed a rigorous framework for studying spatial co-expression of a pair of molecules in the context of spatial transcriptomics (ST) and mass spectrometry imaging (MSI) datasets, at both global (tissue-wide) and local (cell/spot-specific) levels. Existing tools mostly rely on two exploratory geospatial metrics, namely, bivariate Moran’s *I*^108^ and Lee’s *L*^110^, which lead to highly spurious association inference as demonstrated by our simulation studies. Our proposed approach, SpaceBF, builds on the widely used spatially varying coefficients model^136^, effectively capturing spatial autocorrelation and locally varying co-expression patterns. We introduce a spatial GMRF prior inspired by the fused horseshoe^156^, with a global scale that regulates smoothness along graph edges and edge-specific local scales that allow large discontinuities to escape shrinkage. Setting the local scales to unity recovers the standard ICAR prior. We elucidate the prior’s theoretical properties and evaluate its performance against alternative spatial priors. Integrating the prior both within a Gaussian linear regression model and a more complex negative binomial regression framework^135^, SpaceBF is broadly applicable across various analytical contexts and data types.

We conduct a comprehensive evaluation of the proposed method under challenging simulation scenarios, demonstrating its ability to maintain well-controlled type I error rates alongside strong detection power. In three real-world applications, two spatial transcriptomics (ST) datasets and one mass spectrometry imaging (MSI) dataset, the method exhibits robust performance in identifying biologically meaningful molecular interactions, including ligand-receptor (LR) signaling, keratin binding, and peptide co-localization. Notably, the analysis of the cutaneous melanoma sample reveals spatially variable patterns of cell-cell communication, as assessed through LR interactions. The LR pairs exhibit coordinated over- or under-expression within spatially distinct tissue regions (identified from histology) or specific cell types (inferred from transcriptome-wide gene expression). This level of granular understanding may provide critical insights for developing novel, targeted tissue-specific therapies in broader clinical settings^190–192^.

Using the MST as the spatial graph offers several benefits: (i) uniqueness, removing the need to tune additional graph hyperparameters (e.g., GP lengthscales^97^); (ii) reduced computational burden via an exceptionally sparse precision matrix; and (iii) exact Gibbs updates for local horseshoe scales. In our simulations with spatial autocorrelation generated from a Gaussian process with an exponential kernel and varying lengthscales (but a domain-constant slope), the MST performs well, underscoring its robustness. Nonetheless, restricting the spatial structure to a single spanning tree can exclude salient edges^193^, yielding noisier local slope estimates and overly sharp transition boundaries when coefficients vary spatially. In practice, a moderately denser graph, such as a *k*-NN network with a small *k*, often achieves a better trade-off between computational efficiency and appropriate smoothness, as observed in our simulations. A more principled avenue could be to treat the spanning tree as unknown and update it iteratively within the model^194^. While we leverage the `spam` package^195^ for fast sparse Cholesky factorization, overall complexity is graph-structure dependent (e.g., near *O*(*N*) on trees/MSTs and typically around *O*(*N* ^3*/*2^) time for 2D planar/*k*-NN graphs)^196^. As future work, we will pursue MCMC-free, variational-inference-based estimation to improve scalability^197,198^. We have focused on pairwise analyses thus far; extending to joint modeling will follow prior works^199,200^. Although we have primarily used SpaceBF in ST and MSI datasets, it could also be useful in multiplex immunofluorescence (mIF) or imaging mass cytometry datasets where the molecular outcome of interest is generally immune cell types. To study cell type co-localization in such cases, one could split an mIF image into regular grids and count how many cells of two types (*m, m*^′^) fall into each grid. Assuming that the grid centers are the locations *s*_*k*_’s, a spatial graph can be constructed, and the spatial cell counts *X*^*m*^(*s*_*k*_) and *X*^*m*′^ (*s*_*k*_) can be analyzed using our framework. The package is available at https://github.com/sealx017/SpaceBF/.

## 4 Methods

### 4.1 Gaussian and negative binomial regression models

We assume a single sample or image with *n* spots/cells. Let *X*^*m*^(*s*_*k*_) and *X*^*m*′^ (*s*_*k*_) denote the expression of a pair of molecules *m* and *m*^′^, and *C*(*s*_*k*_) be a vector of *p* covariates observed at spot/cell location *s*_*k*_, for *k* ∈ {1, …, *n*}. For example, *C*(*s*_*k*_) can be the cell type indicator or a vector of cell type proportions^164^. We consider the following Gaussian spatially varying coefficients (SVC) model^136^

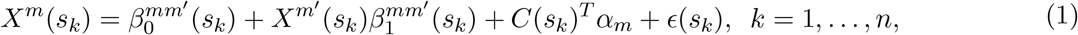

where 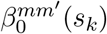 and 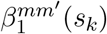 denote spatially varying intercept and slope, respectively, *α*_*m*_ is a fixed effect vector, and *ϵ*(*s*_*k*_) is an independent error term. To interpret the model, a significantly positive 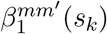 implies that the molecules (*m, m*^′^) co-express at the location *s*_*k*_, while a significantly negative value suggests avoidance. Intuitively, 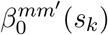 accounts for spatial autocorrelation of molecule *m*. More discussion on the underlying bivariate spatial process is provided in the Supplementary Material. For a count-valued *X*^*m*^(*s*_*k*_) (e.g., genes in the ST datasets), we consider a spatially varying negative binomial (NB) distribution^135^ as *NB*(*ψ*_*m*_(*s*_*k*_), *r*_*m*_) with the failure probability *ψ*_*m*_(*s*_*k*_) modeled as^201^

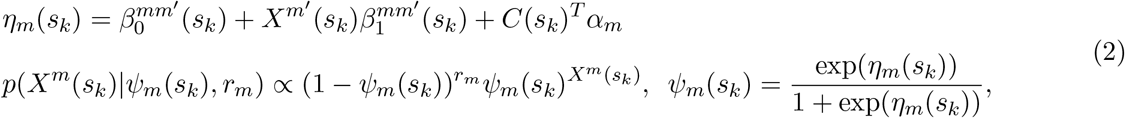

where *p*(.|.) denotes the conditional probability mass function (PMF) and the dispersion parameter *r*_*m*_(*>* 0) is assumed to be constant across locations. To explain how this framework effectively models overdispersion: as *r*_*m*_ → ∞, it reduces to a Poisson model; in contrast, as *r*_*m*_ → 0, the counts become increasingly dispersed relative to the Poisson distribution^135^. Admittedly, this model is limited as the count-valued nature of the molecule *X*^*m*′^ (*s*_*k*_) is not prioritized, appearing as a spatially varying predictor. Jointly modeling (*X*^*m*^(*s*_*k*_), *X*^*m*′^ (*s*_*k*_)) as bivariate NB (BNB) random variables is a possible approach that we do not pursue, as the existing definitions of the BNB distribution (outside of Copula-based constructions)^202–205^ vary considerably, often leading to restrictive correlation structures and inefficient MCMC sampling. To clarify, all applications in the manuscript assume no covariates, i.e., we do not include *C*(*s*_*k*_) or *α*_*m*_, for simplicity. The models in Eqs. 1 and 2 are over-parametrized and do not incorporate spatial dependency between 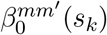 and 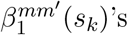, which we discuss next. Let *G* = (*V, E*) denote the MST network between the locations constructed using the *L*_2_ distance for a pair 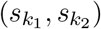, where *V* and *E* are the sets of vertices and edges, respectively. Given a connected, weighted graph, an MST is an acyclic subgraph that connects all vertices and minimizes the sum of the weights of the included edges. Because of this property, MST is routinely used to develop transportation and telecommunication networks^206^. For a regular grid, MST is not unique, as the inter-point distances are not distinct. However, a unique random MST can be curated by simply adding small random values to the distances^207,160^.

### 4.2 Spatial modeling

#### 4.2.1 Spatial fused lasso

In a recent study^157^ of the temperature-salinity relationship in the Atlantic Ocean, Li et al. (2019) consider Eq. 1 and elegantly promote spatial homogeneity of the coefficients by considering fused lasso penalties^151,208^: 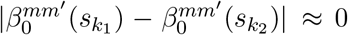 and 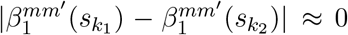 for 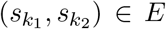, in a frequentist setup. These constraints are intuitive, as it is reasonable to expect both the degree of co-expression, 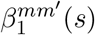, and the effect of “unmeasured” factors, 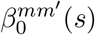, to remain homogeneous across adjacent or connected locations. Extending this idea, a Bayesian fused lasso^209,153^ approach can be considered with Laplacian priors on the pair-wise differences of the coefficients as

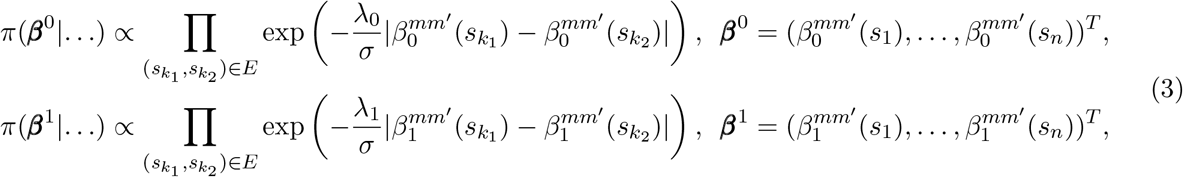

where *σ*^2^ is the variance of the error term *ϵ*(*s*_*k*_), *λ*_0_ and *λ*_1_ are regularization parameters that control the strength of fusion and are assumed to follow gamma priors. Note that *σ*^2^ is only present in the Gaussian model (Eq. 1) and could be omitted from the above exponents for simplicity. We discuss the resemblance of the prior to the intrinsic CAR (ICAR) prior^146^ and, more generally, the intrinsic GMRF (IGMRF) prior^150^ in the Supplementary Material. Theoretically, using the *L*_1_ distance seems appealing, as it has the potential to achieve better spatial smoothing by “exactly” fusing coefficient values at adjacent locations, unlike the *L*_2_ distance implied by the ICAR prior. This is analogous to how lasso regression enforces sparsity in solutions, while ridge regression only shrinks effect sizes toward 0^210^. For transparency, such a spatial fused lasso prior has already been proposed in the existing literature^194,211^.

#### 4.2.2 Spatial fused horseshoe

In variable selection problems, failure of the Bayesian lasso or Laplacian prior to achieve exact sparsity, unlike the frequentist analog, has been reported, while also underestimating larger effect sizes^212–214^. Consequently, the Bayesian fused lasso might struggle to promote spatial smoothness and preserve distinct local features simultaneously. For variable selection, the advantages of the horseshoe prior have been convincingly demonstrated to handle unknown sparsity and large outlying signals^154,215,216^. The horseshoe prior belongs to the class of global-local shrinkage priors^217^, characterized by a “global” hyperparameter that controls overall shrinkage, while “local” hyperparameters control shrinkage per coefficient. Following recent developments on fused horseshoe priors^156,218^, we place horseshoe shrinkage on *edgewise differences* of the spatial coefficients. For a pair of locations 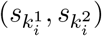 connected by edge *i* ∈ *E*, let 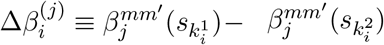 denote the difference, for *j* ∈ {0, 1}. We specify

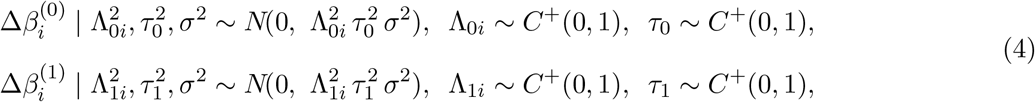

independently across edges *i* = 1, …, *p*, where |*E*|= *p* (e.g., *p* = *N* − 1 for the MST). Let *D* ∈ ℝ^*p*×*N*^ be an oriented incidence matrix and define the weighted Laplacians 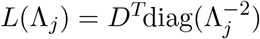 *D* with 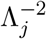 the vector of edgewise precisions 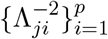. The above construction induces the following intrinsic GMRF prior

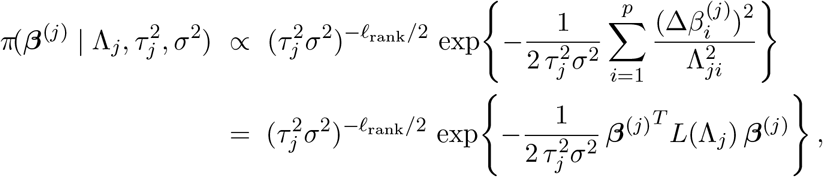

where *ℓ*_rank_ = rank (*L*(Λ_*j*_)) = *N* − *C* where *C* is the number of connected components of the graph (*ℓ*_rank_ = *N* − 1 for a connected graph). When all local scales are set to one, Λ_*ji*_ ≡ 1, the prior reduces to the standard ICAR prior (up to a scale factor)^97^. The deliberately omitted normalizing factor (the generalized determinant of *L*(Λ_*j*_)) depends on Λ_*j*_ but not on ***β***^(*j*)^ or 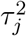; it therefore plays no direct role in the Gibbs updates for ***β***^(*j*)^ or 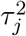. In our implementation, we update the local scales Λ_*ji*_ with conjugate per-edge steps, which are exact on trees and constitute a composite or pseudo-likelihood approximation on general graphs^219–221^. The fused horseshoe’s half-Cauchy global and local scales produce heavy tails (allowing large jumps across edges) and an infinitely tall spike at zero (aggressively shrinking small differences), thereby preserving spatial homogeneity while still accommodating sharp local variations. This connection between locally adaptive fusion and GMRFs was emphasized by Faulkner and Minin (2017)^222^ in a longitudinal context, encouraging fusion between coefficients across time points. In the Supplementary Material, we provide the details of the Gibbs sampling steps, which include the Pólya-Gamma data augmentation strategy^201,223^ for the NB model (Eq. 2).

One crucial aspect that deserves elucidation is the working assumption of independence between edge-wise differences. Specifically, for any two edges 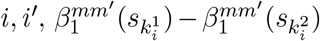 and 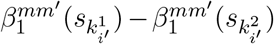 are assumed to be independent in Eq. 4, conditional on the hyperparameters. To see how this assumption could be problematic for a general graph with cycles (i.e., not the MST), we briefly highlight one example from Rue and Held (2005)^150^ provided in the context of IGMRF priors. Suppose there are only three locations *A, B*, and *C*, all neighbors of each other. Letting 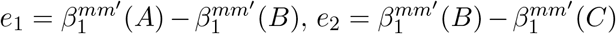, and 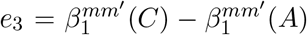, Eq. 4 proceeds to assume *e*_1_, *e*_2_, *e*_3_ are independent and normally distributed with non-identical parameters, yet there is a “hidden” linear constraint, *e*_1_ + *e*_2_ + *e*_3_ = 0, that contradicts independence. Analogously, using a highly connected or dense spatial neighborhood graph *G* introduces numerous hidden constraints corresponding to the cycles in *G*. Interestingly, as shown in Theorem 1 of the Supplementary Material, these constraints need not be enforced explicitly: the posterior sampling distribution of ***β***^0^ and ***β***^1^ under the constraints is unchanged. However, penalizing too many edge-wise differences in the presence of implicit dependencies can lead to oversmoothing and the loss of salient local structure. This consideration naturally favors the MST, which is acyclic (hence no hidden constraints) and removes redundant relationships; moreover, a sparser *G* yields faster Cholesky factorizations of the precision matrix and thus improved computational efficiency. That said, as we show in Section 4.4.3, a *k*-NN graph with a small *k* can significantly outperform the MST in practice.

### 4.3 Hypothesis testing

We consider two types of hypothesis tests: 1) global test: to determine the significance of average association across the entire tissue domain 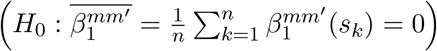, based on the credible interval^224^ of 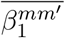, and 2) local test: to determine the significance of location-level association 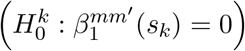, directly based on the credible intervals of 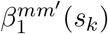’s. Additionally, in the genomic context, having a measure analogous to the frequentist *p*-value is often beneficial. To this end, we utilize a metric termed the probability of direction (*p*_*d*_), which quantifies the probability (between 0.5 and 1) that a parameter has an effect in a specific direction, either positive or negative^225,226^. Mathematically, it is defined as the proportion of the posterior distribution that shares the same sign as the median. *p*_*d*_ resembles a two-sided frequentist *p*-value as *p*_two-sided_ = 2(1 − *p*_*d*_). It is implemented in the *R* package `bayestestR`^225^.

### 4.4 Simulation design

We consider two different simulation designs as outlined below, for assessing the Type I error and power of the model proposed in Eq. 2. The locations at which the variables are simulated are the same as the previously discussed cutaneous melanoma dataset (*n* = 293). We have observed that the results remain unaffected when a randomly generated set of locations or other real data-based sets of locations are used.

#### 4.4.1 Simulation design 1

In the first design, we directly consider the model from Eq. 2 to generate (**X**^*m*^, **X**^*m*′^) based on two steps. First, we generate an NB-distributed RV, **X**^*m*′^, using Gaussian copula^127^, incorporating spatial dependency between the observations via a kernel covariance matrix *H* with an exponential kernel and varying lengthscale (*l*) parameters^114^. Then, based on the simulated **X**^*m*′^, we generate **X**^*m*^ following Eq. 2 with a fixed slope 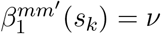. More specifically, for a fixed choice of *l*, failure probability *ψ*_*m*′_, dispersion parameter *r*_*m*′_ for variable *m*^′^, and dispersion parameter *r*_*m*_ for variable *m*, we consider the following steps

1. Simulate a spatially autocorrelated normal RV of size *n* using a Gaussian process (GP) model:

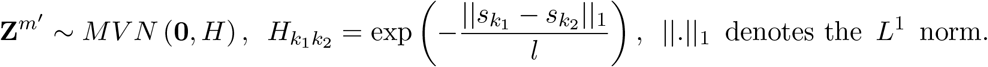
2. Transform to a vector of uniform RVs using the standard normal CDF (Φ):

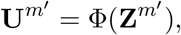
3. Convert to a vector of NB RVs using the inverse CDF of *NB*(*ψ*_*m′*_, *r*_*m′*_), denoted by 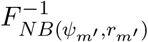:

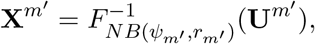

Each element of the resulting vector **X**^*m*′^ retains the marginal NB distribution, *NB*(*ψ*_*m*′_, *r*_*m*′_), where *ψ*_*m*′_ is the failure probability and *r*_*m*′_ is the dispersion.
4. Generate the link function to simulate variable *m* with a fixed slope of 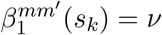 (2):

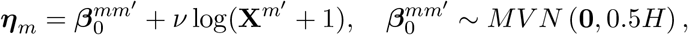
5. Convert the link vector to failure probabilities ***ψ***_*m*_ and simulate **X**^*m*^ from the NB distribution:

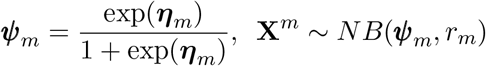

where *r*_*m*_ is a prefixed dispersion parameter. The *k*-th element of **X**^*m*^ follows *NB*(*ψ*_*mk*_, *r*_*m*_), where ***ψ***_*m*_ = (*ψ*_*m*1_, …, *ψ*_*mn*_)^*T*^ and ***η***_*m*_ = (*η*_*m*1_, …, *η*_*mn*_)^*T*^ .

Three values of the lengthscale *l* are considered, *l* = 3.6, 7.2, 18, with the corresponding structure of *H* displayed in Fig. 4A. The failure probability of variable *m*^′^ and dispersion parameters are kept fixed, *ψ*_*m*′_ = 0.5, *r*_*m*_ = *r*_*m*′_ = 1. The slope parameter *ν* is varied between {−0.75, −0.5, −0.25, 0, 0.25, 0.5, 0.75}, with negative and positive values representing negative and positive association, respectively. Higher absolute value of *ν* dictates the strength of association, and *ν* = 0 corresponds to the null model, i.e., **X**^*m*^ and **X**^*m*′^ are independent.

#### 4.4.2 Simulation design 2

In this design, (**X**^*m*^, **X**^*m*′^) are simulated jointly using a bivariate Gaussian copula and spatial dependency incorporated using a bivariate GP framework, where the joint covariance matrix has a Kronecker product structure, comprising a 2 × 2 correlation matrix and the distance kernel covariance matrix *H* (Eq. 9.11 from Banerjee et al. (2014)^97^ and Eq. 6 from the Supplementary Material). Specifically, we consider the following steps

1. Simulate spatially cross-correlated normal RVs:

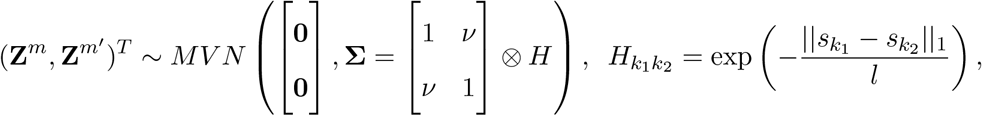
2. Transform to uniform RVs using the standard normal CDF (Φ):

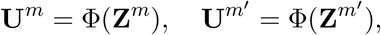
3. Convert to NB random variables using the inverse CDFs:

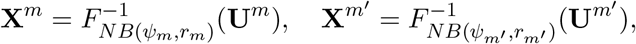

Each element of the resulting vectors **X**^*m*^ and **X**^*m*′^ retain the marginal NB distributions, *NB*(*ψ*_*m*_, *r*_*m*_) and *NB*(*ψ*_*m*′_, *r*_*m*′_), respectively, where *ψ*_*m*_, *ψ*_*m*′_ are failure probabilities and *r*_*m*_, *r*_*m*′_ are dispersions.

The lengthscale *l* is varied between {0.6, 1.8, 3.6, 7.2}. The failure probabilities and dispersion parameters are kept fixed, *ψ*_*m*_ = *ψ*_*m*′_ = 0.5, *r*_*m*_ = *r*_*m*′_ = 1. The parameter *ν* is varied between {−0.75, −0.5, −0.25, 0, 0.25, 0.5, 0.75}, with similar implications on the direction and strength of association as before.

#### 4.4.3 Simulation design 3

We consider the model from Eq. 2 to generate (**X**^*m*^, **X**^*m*′^) based on two steps. First, we generate an NB-distributed RV, **X**^*m*′^, using Gaussian copula^127^, incorporating spatial autocorrelation via *H* with a large lengthscale, *l* = 7.2. Then, based on the simulated **X**^*m*′^, we generate **X**^*m*^ following Eq. 2, this time with a spatially varying slope 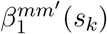. Specifically, we consider the following steps

1. Simulate **X**^*m*′^ following steps 1, 2, and 3 from Section 4.4.1.
2. Simulate the slope 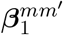 in two ways: based on a) linear partition boundary and b) circular boundary. Let 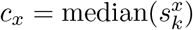, 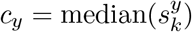 denote the median of *xy*-coordinates, respectively.
  a. *Linear partition boundary:* Define two partitioning subsets of the spatial domain as

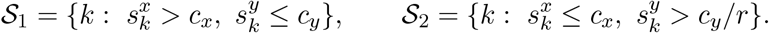

Draw *σ*_*k*_ ∼ Unif(0.3, 0.6). For *k* ∈ 𝒮_1_, draw 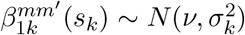; for *k* ∈ 𝒮_2_, draw 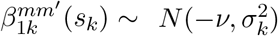; for other *k*’s 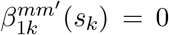. Vary *r* ∈ {1, 2} to create two different partitioning configurations.
  b. *Circular boundary* : Define the indicator for being inside a circle with radius *r* ∈ {1, 2} and centered at the median

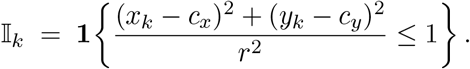

Set *σ*_*k*_ = *σ*_in_ 𝕀_*k*_ + *σ*_out_ (1 − 𝕀_*k*_) with *σ*_in_ = 0.3, *σ*_out_ = 0.6, and draw 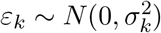. Finally, define

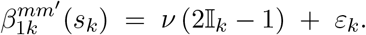

Thus, 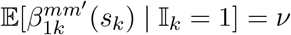 (inside the circle) and 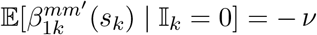 (outside).
3. Generate the link function to simulate variable *m* with the slope 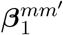 :

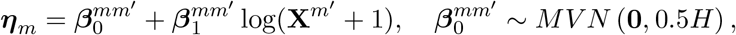
4. Convert the link vector to failure probabilities ***ψ***_*m*_ and simulate **X**^*m*^ from the NB distribution:

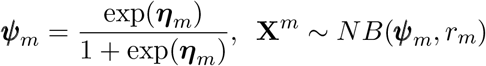

where *r*_*m*_ is a prefixed dispersion parameter. The *k*-th element of **X**^*m*^ follows *NB*(*ψ*_*mk*_, *r*_*m*_), where ***ψ***_*m*_ = (*ψ*_*m*1_, …, *ψ*_*mn*_)^*T*^ and ***η***_*m*_ = (*η*_*m*1_, …, *η*_*mn*_)^*T*^ . Each element of the resulting vectors **X**^*m*^ and **X**^*m*′^ retain the marginal NB distributions, *NB*(*ψ*_*m*_, *r*_*m*_) and *NB*(*ψ*_*m*′_, *r*_*m*′_), respectively, where *ψ*_*m*_, *ψ*_*m*′_ are failure probabilities and *r*_*m*_, *r*_*m*′_ are dispersions.

The dispersion parameters are kept fixed, *r*_*m*_ = *r*_*m*′_ = 1, and the failure probability *ψ*_*m*′_ = 1. The boundary parameter (or radius) *r* is varied between {1, 2}. The boundary (radius) parameter *r* varies over {1, 2}, and the effect-size parameter *ν* varies over {2, 4}. Results for *ν* = 2 are provided in the Supplementary Material, while *ν* = 4 results are shown in Figs. 7 and 8.

**Figure 8:**
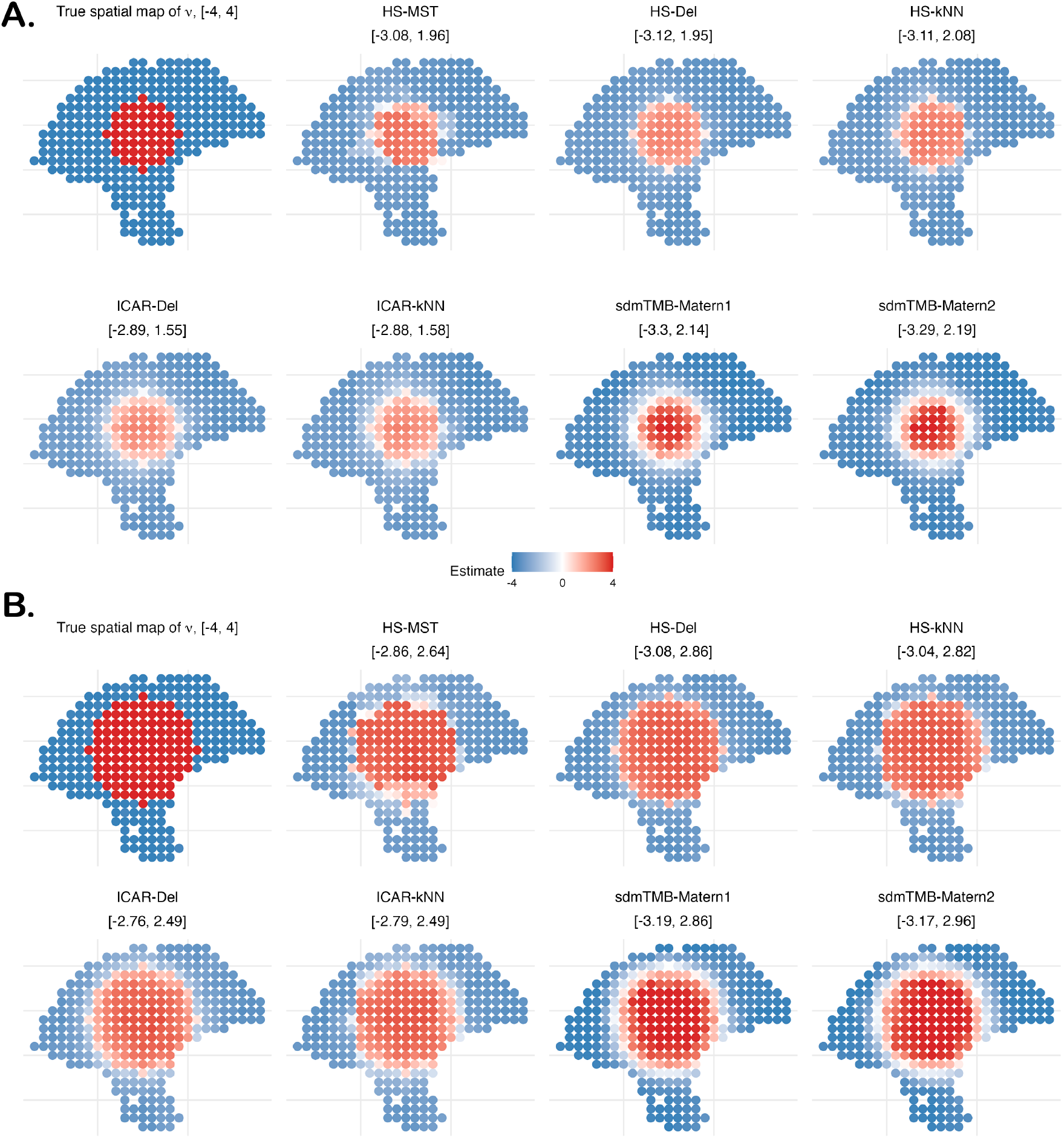
SVC simulation based on circular boundary from Section 4.4.3 with effect size *ν* = 4. **A**. A smaller circle with positive *ν* values, radius *r* = 4, and **B**. A larger circle with positive *ν* values, *r* = 6.

### 4.5 Competing methods

We compare **SpaceBF** against five methods: (1) MERINGUE^98^ with a Delaunay-triangulation graph; (2) SpatialDM^102^ using a Gaussian kernel weight matrix (lengthscale 1.2, as in their original melanoma analysis); (3) Lee’s *L*^110^ on an *ϵ*-neighborhood network (implemented via the *R* package spdep^227^), where *ϵ* is set to the maximum nearest-neighbor distance to ensure connectivity; (4) SpaGene^100^; and (5) PearsonCorr, the standard Pearson correlation. We do not include LIANA+^104^ or Voyager^117^, as the former essentially wraps SpatialDM and the latter directly applies Lee’s *L*. We adopt the *ϵ*-neighborhood instead of a fixed *k*-NN to prevent oversmoothing in Lee’s *L*, while preserving graph connectivity; however, the approach proved futile, as seen from the simulation studies. Table 1 summarizes the methods in terms of their assumptions and limitations. We also attempted to evaluate the performance of SpatialCorr^178^ and Copulacci^105^. SpatialCorr was straightforward to use, but it proved highly sensitive to the choice of the lengthscale *l* in its innovative use of the spatial covariance matrix *H*. Copulacci was slightly difficult to use and will be benchmarked in a future study.

**Table 1:** Comparison of the methods in terms of the underlying assumptions. ^∗^methods that are not evaluated in the simulations.

| Method | Central metric or concept | Global and local co-expression tests | Test type | Potential sensitivity |
| --- | --- | --- | --- | --- |
| **SpaceBF** | Spatially varying coefficients model with NB distribution | Global and local | Exact | Spatial adjacency graph |
| MERINGUE | Bivariate Moran's $I$ | Global | Permutation test | Spatial adjacency graph |
| SpatialDM | Bivariate Moran's $I$ | Global and local | Permutation or exact test | Lengthscale in the kernel weight matrix |
| LIANA+ | Bivariate Moran's $I$ and cosine similarity | Global and local | Permutation test | As above |
| Voyager | Lee's $L$ | Global and local | Permutation test | Spatial adjacency graph |
| SpaGene | $k$-NN network and earth mover's distance | Global but with local interaction scores | Permutation test | Choice of $k$ |
| PearsonCorr | Pearson correlation | Global | Permutation or exact test | None |
| SpatialCorr * | Spatial kernel-weighted sample correlation | Global and local | Permutation test | Lengthscale in the kernel weight matrix |
| Copulacci * | Bivariate Poisson distribution along an adjacency graph | Global but with local interaction scores | Permutation test | Spatial adjacency graph and fixed correlation term across locations |

We further evaluate the performance of SpaceBF (the spatial horseshoe) on general spatial graphs, alongside standard spatial priors, within our SVC framework. The configurations (methods shown in Figs. 7 and 8) considered are summarized in Table 2, with brief notes on their limitations.

**Table 2:** Methods, graphs, priors, and limitations. HS can be placed on any spatial adjacency graph; denser graphs (e.g., Delaunay or large-*k k*NN) increase the risk of over-smoothing. HS-Del and HS-*k*NN (*k*=3) probe denser and intermediate backbones than HS-MST. ICAR variants are included for comparison. The `sdmTMB` Matérn models differ by mesh density; a coarser mesh leads to a smoother spatial field.

| Method | Graph | Prior | Limitations |
| --- | --- | --- | --- |
| HS-MST | Minimum spanning tree (MST) | Horseshoe (HS) | May under-smooth within dense micro-domains |
| HS-Del | Delaunay triangulation (Del) | HS | Risk of over-smoothing and spurious association; approximate local scale update |
| HS-$k$NN | $k$-nearest neighbors ($k=3$) | HS | Sensitive to $k$: larger $k$ increases density and risk of over-smoothing |
| ICAR-Del | Delaunay triangulation | ICAR (special case of HS) | Risk of diffusing boundaries on dense graphs; tends to attenuate sharp slope changes. |
| ICAR-$k$NN | $k$-nearest neighbors | ICAR | Similar boundary diffusion; performance depends on $k$ |
| sdmTMB-Matérn1 | Mesh (cutoff = 1; denser) | Matérn (SPDE) | Risk of diffusing boundaries; mesh tuning adds burden, Laplace approximation may not converge |
| sdmTMB-Matérn2 | Mesh (cutoff = 1.5; coarser) | Matérn (SPDE) | Above problems and risk of oversmoothing |

## 5 Runtime comparison and convergence diagnostics

In most analyses we ran 5,000 MCMC iterations with 2,500 burn-in. We compared runtimes for our package `SpaceBF` across priors and spatial backbones (from sparser to denser). Figure 9 shows that HS and ICAR have comparable runtimes, scaling approximately linearly with *n*. Denser graphs (e.g., *k*-NN with *k* = 9) are modestly slower. For *n* = 5,000, `SpaceBF` completes in about 12 minutes on a Mac Pro (M3 Max). For substantially larger datasets, a practical alternative is `sdmTMB`^162^ for NB SVC via Laplace-approximate maximum likelihood: it is extremely fast but can be less precise, may fail to converge, and often requires tuning the mesh density.

**Figure 9:**
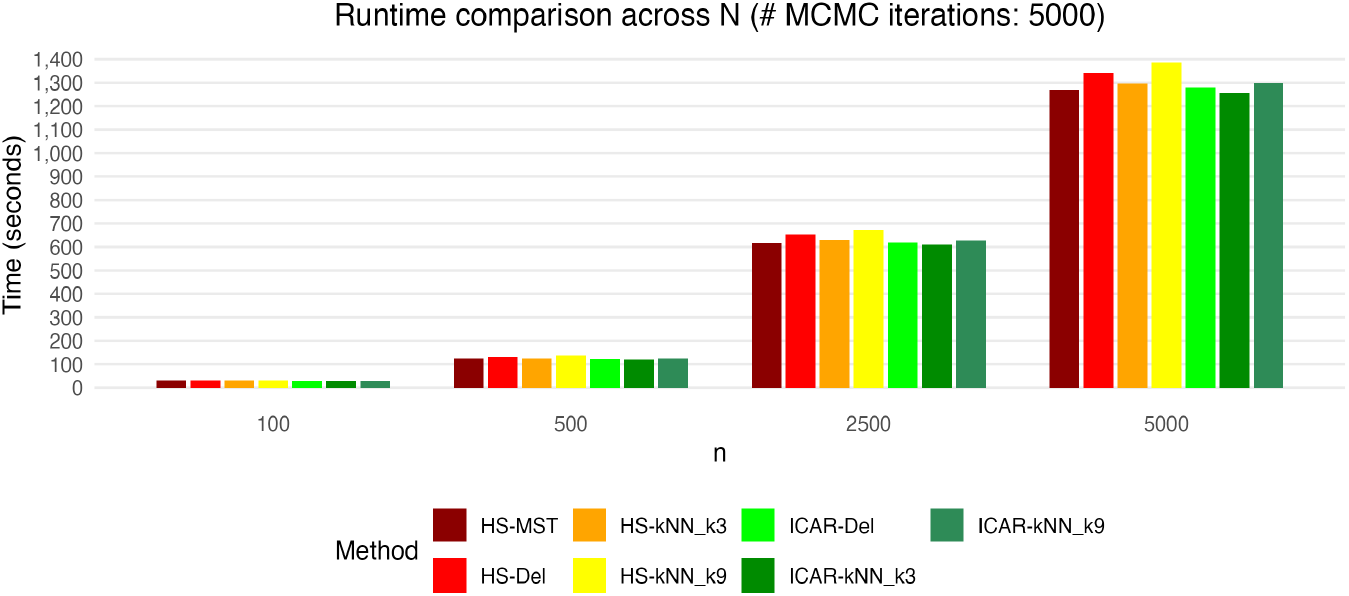
Run-time comparison of `SpaceBF`, with different priors: the horseshoe GMRF and ICAR, with varying spatial adjacency graphs.

For the convergence diagnostics, we computed the Geweke statistic^228^ for each 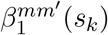, implemented in the *R* package *coda*^229^, and investigated the trace plots of a few randomly chosen 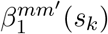 ‘s (see the Supplementary Material). When either the variable *m* or *m*^′^ is highly sparse (*>* 75% zeroes), imposing additional normal priors on 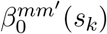 ‘s and 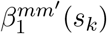 ‘s with a moderate variance, such as *N* (0, 10), drastically improves mixing and overall convergence performance.

## 6 Data and software availability

The melanoma and cSCC datasets are publicly available at the links: 1) cutaneous melanoma^39^: https://zenodo.org/records/8215682 with sample ID “`ST_mel1_rep2`”, and 2) cSCC^163^: https://www.ncbi.nlm.nih.gov/geo/query/acc.cgi?acc=GSE144240 with sample ID “`GSM4284236 P6_cSCC_scRNA`”. The spatial proteomics dataset is available on Zenodo at https://zenodo.org/records/15866928. A GitHub *R* package named `SpaceBF`, with the proposed method and the first two datasets in “.rda” format, is available at https://github.com/sealx017/SpaceBF/.

## 8 Acknowledgments

The authors thank Dr. Peggi Angel from the Medical University of South Carolina for her help in the spatial proteomics dataset acquisition and interpretation. S.S. and B.N. contributed equally to the conceptualization and methodology of the project, and jointly wrote the first draft. S.S. conducted the validation, simulation experiments, and software development. The authors do not have any competing interests.

