## Supplementary Material for "SpaceBF: Spatial coexpression analysis using Bayesian Fused approaches in spatial omics datasets"

Souvik Seal and Brian Neelon

Department of Public Health Sciences, College of Medicine, Medical University of South Carolina, Charleston, USA

June 2025

### 1 A brief review of existing spatial priors

In the Bayesian framework, the spatial structure of the parameter vectors can be modeled in several ways:

1. *Gaussian process (GP) model*: We can assume that $\beta_0^{mm'}(s)$ and $\beta_0^{mm'}(s)$ follow stationary zero-centered Gaussian spatial process models [1] with covariance functions: $\text{cov}(\beta_0^{mm'}(s), \beta_0^{mm'}(s')) = \sigma_0^2\rho_0(||s - s'||, \phi_0)$ and $\text{cov}(\beta_1^{mm'}(s), \beta_1^{mm'}(s')) = \sigma_1^2\rho_1(||s - s'||, \phi_1)$, where $\rho_0, \rho_1$ are valid correlation functions with hyperparameters $\phi_1$, $\phi_2$, and $\sigma_0^2$, $\sigma_1^2$ are variances. The joint distribution of $\boldsymbol{\beta}_0^{mm'} = (\beta_0^{mm'}(s_1), \ldots, \beta_0^{mm'}(s_n))^T$ and $\boldsymbol{\beta}_1^{mm'} = (\beta_1^{mm'}(s_1), \ldots, \beta_0^{mm'}(s_n))^T$ can be written as
$$\boldsymbol{\beta}_0^{mm'} \sim MVN(0, \sigma_0^2 H_0(\phi_0)), \;\; [[H_0(\phi_0)]]_{kk'} = \rho_0(||s_k - s_{k'}||, \phi_0)$$
$$\boldsymbol{\beta}_1^{mm'} \sim MVN(0, \sigma_1^2 H_1(\phi_1)), \;\; [[H_1(\phi_1)]]_{kk'} = \rho_1(||s_k - s_{k'}||, \phi_1).$$
2. *Conditionally autoregressive (CAR) model*: Let $W = [[W_{kk'}]]$ be a binary proximity matrix between $n$ locations, with $W_{kk'} = 1$ if locations $s_k$ and $s_{k'}$ fall within a preset distance or 0 otherwise ($k \neq k'$).

Let $W_{k+} = \sum_{k'} W_{kk'}$ be the $k$-th row-sum of $W$, and $D_w = diag(W_{k+})$. Univariate CAR priors on the two sets of coefficients can be written as

$$\beta_0^{mm'}(s_k)|\beta_0^{mm'}(s'_k), k' \neq k \sim N\left(p_0 \sum_{k'} \frac{W_{kk'}}{W_{k+}} \beta_0^{mm'}(s'_k), \sigma_0^2 \frac{1}{W_{k+}}\right),$$
$$\beta_1^{mm'}(s_k)|\beta_1^{mm'}(s'_k), k' \neq k \sim N\left(p_1 \sum_{k'} \frac{W_{kk'}}{W_{k+}} \beta_1^{mm'}(s'_k), \sigma_1^2 \frac{1}{W_{k+}}\right), \quad k = 1, \ldots, n.$$

where $p_0$ and $p_1$ are "propriety" constants, both less than 1 and such that $(D_w - p_0 W)$ and $(D_w - p_1 W)$ are positive definite (PD). Using matrix notation, the distributions can be written as

$$\pi(\boldsymbol{\beta}_0^{mm'}|.) \propto \exp\left(-\frac{1}{2\sigma_0^2}\boldsymbol{\beta}_0^{mm',T}(D_w - p_0 W)\boldsymbol{\beta}_0^{mm'}\right),$$
$$\pi(\boldsymbol{\beta}_1^{mm'}|.) \propto \exp\left(-\frac{1}{2\sigma_1^2}\boldsymbol{\beta}_1^{mm',T}(D_w - p_1 W)\boldsymbol{\beta}_1^{mm'}\right).$$

$p_0 = p_1 = 1$ leads to a special class of priors known as intrinsic CAR (ICAR) prior. Since an ICAR prior distribution is improper (resulting posteriors are proper), a small diagonal adjustment to the precision matrix $(D_w - W)$ might improve performance [2, 3]. Unlike a GP prior that assumes an unobserved infinite process to model the continuous spatial dependence, the CAR and ICAR priors use a spatially defined graph structure, assuming a finite process on a discrete set of points.

3. *Gaussian Markov random field (GMRF)*: Let $G = (V, E)$ be an undirected graph between the points constructed based on the $L^2$ distance, where $V$ denotes the set of vertices, i.e., $\{s_1, \ldots, s_n\}$, and $E$ denotes the set of edges. Assume that an edge $\{s_k, s_{k'}\}$ is absent in $E$ if and only if the node values (coefficient values) at locations $s_k$ and $s_{k'}$ are conditionally independent given the remaining node values. Then the GMRF priors based on $G$ posit the following distributions

$$\pi(\boldsymbol{\beta}_0^{mm'}|.) \propto \exp\left(-\frac{1}{2\sigma_0^2}\boldsymbol{\beta}_0^{mm',T} Q_0 \boldsymbol{\beta}_0^{mm'}\right) \text{ s.t. } [[Q_0]]_{kk'} \neq 0 \text{ iff } \{s_k, s_{k'}\} \in E,$$
$$\pi(\boldsymbol{\beta}_1^{mm'}|.) \propto \exp\left(-\frac{1}{2\sigma_1^2}\boldsymbol{\beta}_1^{mm',T} Q_1 \boldsymbol{\beta}_1^{mm'}\right) \text{ s.t. } [[Q_1]]_{kk'} \neq 0 \text{ iff } \{s_k, s_{k'}\} \in E$$

where $Q_0$ and $Q_1$ are precision matrices, which incorporate the conditional dependence or connectivity in the network $G$. The precision matrices are allowed to be singular. It can be observed that the CAR prior is a special type of GMRF prior. In fact, *the pairwise difference GMRF prior* [4] leads to the following densities

$$
\begin{aligned}
\pi(\boldsymbol{\beta}_0^{mm'}|.) &\propto \exp\left(-\frac{1}{2\sigma_0^2}\sum_{(s_k,s_{k'})\in E}(\beta_0^{mm',T}(s_k)-\beta_0^{mm',T}(s_{k'})\right)^2, \\
\pi(\boldsymbol{\beta}_1^{mm'}|.) &\propto \exp\left(-\frac{1}{2\sigma_1^2}\sum_{(s_k,s_{k'})\in E}(\beta_1^{mm',T}(s_k)-\beta_1^{mm',T}(s_{k'})\right)^2
\end{aligned} \tag{1}
$$

which are the same as ICAR priors [1]. Of note, replacing the $L^2$ distance with the $L^1$ distance or any general even function $\Phi(z)$, increasing w.r.t. $|z|$, is briefly discussed in Besag (1991) [5].

In all the above models, we have ignored the correlation between $\boldsymbol{\beta}_0^{mm'}$ and $\boldsymbol{\beta}_1^{mm'}$, which is typically modeled using a Kronecker-product structure. The GP models are computationally expensive (because the hyperparameters $\phi_0, \phi_1$ need to be tuned) and require further approximations such as predictive process [6] and nearest neighbor Gaussian process [7] to be applicable on high-dimensional datasets. The GMRF and CAR priors are computationally more tractable, with further potential for scalability through the use of advanced approximation techniques, such as the integrated nested Laplace approximation (INLA) [4].

### 2 More discussions on the proposed approach

Let $G = (V, E)$ denote the MST network between the locations constructed using the $L^2$ distance, where $V$ and $E$ are the sets of vertices and edges, respectively.

#### 2.1 Spatial fused lasso and its connection to GMRF

Following the main text, the Laplacian priors on the pair-wise differences of coefficients can be written as

$$
\begin{aligned}
\pi(\boldsymbol{\beta}_0^{mm'}|\ldots) &\propto \prod_{(s_{k_1},s_{k_2})\in E}\exp\left(-\frac{\lambda_0}{\sigma}|\beta_0^{mm'}(s_{k_1})-\beta_0^{mm'}(s_{k_2})|\right),\ \boldsymbol{\beta}_0^{mm'}=(\beta_0^{mm'}(s_1),\ldots,\beta_0^{mm'}(s_n))^T, \\
\pi(\boldsymbol{\beta}_1^{mm'}|\ldots) &\propto \prod_{(s_{k_1},s_{k_2})\in E}\exp\left(-\frac{\lambda_1}{\sigma}|\beta_1^{mm'}(s_{k_1})-\beta_1^{mm'}(s_{k_2})|\right),\ \boldsymbol{\beta}_1^{mm'}=(\beta_1^{mm'}(s_1),\ldots,\beta_1^{mm'}(s_n))^T,
\end{aligned} \tag{2}
$$

where $\sigma^2$ is the variance of the error term $\epsilon(s_k)$, present only in the Gaussian model. Scaling by $\sigma$ inside the exponent in Eq. 2 is motivated by Park and Casella (2008) [8], who showed its necessity to ensure a globally optimal solution in the context of a simple Bayesian lasso. However, in our simulations, the effect was minimal. In theory, using an $L_1$ distance shall be more effective in "exactly" fusing the coefficient values at two spatially close locations, compared to an $L_2$ distance. As mentioned earlier, in the seminal paper by Besag (see Eq. 4.4 of Besag (1991) [5]), such a prior is recommended "if discontinuities in the risk surface are expected.".

For the posterior sampling, the Laplacian likelihood can be expanded as a superposition of an infinite number of Gaussian distributions [9],

$$\frac{\sqrt{\lambda}}{2}\exp\Big[-\sqrt{\lambda}|x|\Big] = \int_0^{\infty}\sqrt{\frac{1}{2\pi\zeta}}\exp\Big[-\frac{x^2}{2\zeta}\Big]\frac{\lambda}{2}\exp\Big[-\frac{\lambda\zeta}{2}\Big]\,d\zeta$$

Suppose there are $p$ edges in $E$. Exploiting the above property, we introduce two new latent mixing vectors of length $p$ as $\boldsymbol{\zeta}_0 = (\zeta_{10},\ldots,\zeta_{p0})^T$ and $\boldsymbol{\zeta}_1 = (\zeta_{11},\ldots,\zeta_{p1})^T$, where $\zeta_{i0}$ and $\zeta_{i1}$ correspond to $|\beta_0^{mm'}(s_{k_i^1}) - \beta_0^{mm'}(s_{k_i^2})|$ and $|\beta_1^{mm'}(s_{k_i^1}) - \beta_1^{mm'}(s_{k_i^2})|$, respectively, and $(k_i^1, k_i^2) \in \{1,\ldots,n\}$ are the nodes associated with the $i$-th edge from $E$. We then write the conditional prior of $(\boldsymbol{\beta}_0^{mm'}, \boldsymbol{\beta}_1^{mm'})$ as

$$\begin{aligned}
\pi(\boldsymbol{\beta}_0^{mm'}|\boldsymbol{\zeta}_0,\sigma^2,.) &= \prod_{i=1}^{p}\sqrt{\frac{1}{2\pi\zeta_{i0}\sigma^2}}\exp\left[-\frac{\left(\beta_0^{mm'}(s_{k_i^1}) - \beta_0^{mm'}(s_{k_i^2})\right)^2}{2\zeta_{i0}\sigma^2}\right]\\
\pi(\boldsymbol{\beta}_1^{mm'}|\boldsymbol{\zeta}_1,\sigma^2,.) &= \prod_{i=1}^{p}\sqrt{\frac{1}{2\pi\zeta_{i1}\sigma^2}}\exp\left[-\frac{\left(\beta_1^{mm'}(s_{k_i^1}) - \beta_1^{mm'}(s_{k_i^2})\right)^2}{2\zeta_{i1}\sigma^2}\right]
\end{aligned} \tag{3}$$

The mixture components follow the joint distribution $\pi(\boldsymbol{\zeta}_0,\boldsymbol{\zeta}_1) = \prod_{i=1}^{p}\Big[\frac{\lambda_0}{2}\frac{\lambda_1}{2}\exp\Big[-\frac{\lambda_0\zeta_{i0}}{2} - \frac{\lambda_1\zeta_{i1}}{2}\Big]\Big]$. We notice that Eq. 3 is a weighted version of the pairwise difference GMRF prior or ICAR prior from Eq. 1.

### 2.2 MCMC scheme for both priors in the Gaussian model

For notational simplicity, we drop the covariates vector $Z(s_k)$ and the associated coefficient vector $\alpha_m$ in the following derivations. It is straightforward to include these into the framework.

#### 2.2.1 Spatial fused lasso

We multiply the densities from Eq. 3 and simplify using a matrix notation as

$$\pi(\boldsymbol{\beta}^{mm'}|.) \propto \left[\frac{1}{\prod_{i=1}^{p}\sqrt{\zeta_{i0}\zeta_{i1}}}\right] \exp\left[-\frac{1}{2\sigma^2}(\boldsymbol{\beta}^{mm'})^T B \boldsymbol{\beta}^{mm'}\right] \tag{4}$$

where $\boldsymbol{\beta}^{mm'} = (\boldsymbol{\beta}_0^{mm'}, \boldsymbol{\beta}_1^{mm'})^T$ and $B$ is a block-diagonal matrix with two blocks $B_0$ and $B_1$ defined as $[B_0]_{k_i^1,k_i^2} = [B_0]_{k_i^2,k_i^1} = -\frac{1}{\zeta_{i0}}$, $[B_1]_{k_i^1,k_i^2} = [B_1]_{k_i^2,k_i^1} = -\frac{1}{\zeta_{i1}}$, $i = 1,\dots,p$, (and 0 elsewhere in the off-diagonal), $[B_0]_{k,k} = -\sum_{k'\neq k}[B_0]_{k,k'}$, $[B_1]_{k,k} = -\sum_{k'\neq k}[B_1]_{k,k'}$.

If $\sigma^2$ is omitted from Eqs. 3 and 4, we can redefine $B$ as $\sigma^2 B$ and keep the subsequent derivation as it is. Assuming $\epsilon(s_k)$'s follow IID $N(0,\sigma^2)$, we write the distribution of $\mathbf{X}^m$, without the priors, as

$$p(\mathbf{X}^m|\mathbf{X}^{*m'}, \boldsymbol{\beta}^{mm'}, \sigma^2) \propto \exp\left[-\frac{1}{2\sigma^2}((\mathbf{X}^m - \mathbf{X}^{*m'}\boldsymbol{\beta}^{mm'})^T(\mathbf{X}^m - \mathbf{X}^{*m'}\boldsymbol{\beta}^{mm'})\right], \tag{5}$$

where $\mathbf{X}^{*m'} = [I_n,\ diag(\mathbf{X}^{m'})]_{n\times 2n}$, where $I_n$ is the identity matrix of dimension $n$. Now, the required conditional posterior distributions for a Gibbs sampling are derived as

1. $$p(\boldsymbol{\beta}^{mm'}|.) \propto \exp\left[-\frac{1}{2\sigma^2}(\boldsymbol{\beta}^{mm'} - M)^T\Sigma^{-1}(\boldsymbol{\beta}^{mm'} - M)\right],$$
$$\Sigma = \left(B + (\mathbf{X}^{*m'})^T\mathbf{X}^{*m'}\right)^{-1},\ \ M = \Sigma\left((\mathbf{X}^{*m'})^T\mathbf{X}^m\right)$$

2. $$p(\zeta_{i0}|.) \propto \zeta_{i0}^{-1/2}\exp\left[-\frac{1}{2}\left(\frac{\left(\beta_0^{mm'}(s_{k_i^1}) - \beta_0^{mm'}(s_{k_i^2})\right)^2}{\zeta_{i0}\sigma^2} + \lambda_0\zeta_{i0}\right)\right]$$
$$p(\zeta_{i1}|.) \propto \zeta_{i1}^{-1/2}\exp\left[-\frac{1}{2}\left(\frac{\left(\beta_1^{mm'}(s_{k_i^1}) - \beta_1^{mm'}(s_{k_i^2})\right)^2}{\zeta_{i1}\sigma^2} + \lambda_1\zeta_{i1}\right)\right]$$

3. Assuming an improper prior on $\sigma^2$: $\pi(\sigma^2) = 1/\sigma^2$,
$$p(\sigma^2|.) \propto \left(\frac{1}{\sigma^2}\right)^{n/2+(2n)/2+1}\exp\left[-\frac{1}{2\sigma^2}((\mathbf{X}^m - \mathbf{X}^{*m'}\boldsymbol{\beta}^{mm'})^T(\mathbf{X}^m - \mathbf{X}^{*m'}\boldsymbol{\beta}^{mm'}) + (\boldsymbol{\beta}^{mm'})^T B\boldsymbol{\beta}^{mm'})\right]$$

4. Putting a gamma prior with hyper-parameters $\delta_{1s}, \delta_{2s}$ on the shrinkage parameter $\lambda_s$, the corresponding posterior distribution can be derived as
$$\pi(\lambda_s|\delta_{1s},\delta_{2s}) = \frac{\delta_{1s}^{\delta_{2s}}}{\Gamma(\delta_{2s})}\lambda_s^{\delta_{2s}-1}\exp(-\delta_{1s}\lambda_s),\ \text{ for }\ s = 0, 1,$$
$$p(\lambda_s|.) \propto \left[\lambda_s^{p+\delta_{2s}-1}\right]\exp\left[-\lambda_s\left(\delta_{1s} + \frac{1}{2}\sum_{i=1}^{p}\zeta_{is}\right)\right],$$

##### 2.2.2 Spatial fused horseshoe

To derive the Gibbs sampling algorithm for spatial fused horseshoe, we introduce new latent variables, $\gamma_{0i}, \gamma_{1i}, \epsilon_0, \epsilon_1$ to write the half-Cauchy priors as mixtures of inverse gamma ($IG$) priors [10] as below

$$
\begin{aligned}
\Lambda_{0i}^2|\gamma_{0i} &\sim IG\left(\frac{1}{2}, \frac{1}{\gamma_{0i}}\right), & \Lambda_{1i}^2|\gamma_{1i} &\sim IG\left(\frac{1}{2}, \frac{1}{\gamma_{1i}}\right),\\
\gamma_{0i} &\sim IG\left(\frac{1}{2}, 1\right), & \gamma_{1i} &\sim IG\left(\frac{1}{2}, 1\right),\\
\tau_0^2|\epsilon_0 &\sim IG\left(\frac{1}{2}, \frac{1}{\epsilon_0}\right), & \tau_1^2|\epsilon_1 &\sim IG\left(\frac{1}{2}, \frac{1}{\epsilon_1}\right),\\
\epsilon_0 &\sim IG\left(\frac{1}{2}, 1\right), & \epsilon_1 &\sim IG\left(\frac{1}{2}, 1\right).
\end{aligned}
$$

We redefine the matrix $B$ from the previous section as

$$
\begin{aligned}
&[B_0]_{k_i^1,k_i^2} = [B_0]_{k_i^2,k_i^1} = -\frac{1}{\tau_0^2\Lambda_{0i}^2}, \ \ [B_1]_{k_i^1,k_i^2} = [B_1]_{k_i^2,k_i^1} = -\frac{1}{\tau_1^2\Lambda_{1i}^2},\\
&[B_0]_{k,k} = -\sum_{k'\neq k}[B_0]_{k,k'}, \ \ [B_1]_{k,k} = -\sum_{k'\neq k}[B_1]_{k,k'}.
\end{aligned}
$$

The required conditional posterior distributions for a Gibbs sampling will then be

1. Posterior distributions highlighted in points 1-3 from section 2.2.1.

2. $$
\begin{aligned}
&\Lambda_{0i}^2|. \sim IG\left(1, \frac{1}{\gamma_{0i}} + \frac{\left(\beta_0^{mm'}(s_{k_i^1}) - \beta_0^{mm'}(s_{k_i^2})\right)^2}{2\tau_0^2\sigma^2}\right), \ \ \gamma_{0i}|. \sim IG\left(1, 1 + \frac{1}{\Lambda_{0i}^2}\right),\\
&\tau_0^2|. \sim IG\left(\frac{p+1}{2}, \frac{1}{\epsilon_0} + \sum_{i=1}^{p}\frac{\left(\beta_0^{mm'}(s_{k_i^1}) - \beta_0^{mm'}(s_{k_i^2})\right)^2}{2\Lambda_{0i}^2\sigma^2}\right), \ \ \epsilon_0|. \sim IG\left(1, 1 + \frac{1}{\tau_0^2}\right),\\
&\Lambda_{1i}^2|. \sim IG\left(1, \frac{1}{\gamma_{1i}} + \frac{\left(\beta_1^{mm'}(s_{k_i^1}) - \beta_1^{mm'}(s_{k_i^2})\right)^2}{2\tau_1^2\sigma^2}\right), \ \ \gamma_{1i}|. \sim IG\left(1, 1 + \frac{1}{\Lambda_{1i}^2}\right),\\
&\tau_1^2|. \sim IG\left(\frac{p+1}{2}, \frac{1}{\epsilon_1} + \sum_{i=1}^{p}\frac{\left(\beta_1^{mm'}(s_{k_i^1}) - \beta_1^{mm'}(s_{k_i^2})\right)^2}{2\Lambda_{1i}^2\sigma^2}\right), \ \ \epsilon_1|. \sim IG\left(1, 1 + \frac{1}{\tau_1^2}\right).
\end{aligned}
$$

### 2.3 MCMC scheme for an NB model using Pólya-gamma augmentation

For the NB model, we use a data-augmented Gibbs sampler proposed by Pillow and Scott (2012) [11] and Polson et al. (2013) [12] for posterior sampling derivations. The sampler introduces latent Pólya-Gamma ($PG$)-distributed weights $w_k$, for $k = 1, \ldots, n$. For $b > 0$ and $c \in \mathbb{R}$, a random variable $M$ is said to have a $PG$ distribution if

$$M \sim PG(b, c) = \frac{1}{2\pi^2} \sum_{g=1}^{\infty} \frac{d_g}{(g - 1/2)^2 + c^2/(4\pi^2)}$$

where $d_g$'s are independent $Gamma(b, 1)$ random variables. The NB mass function (Eq. 2 from the main text) can be expanded as a mixture of distributions as

$$p(X^m(s_k)|\psi_m(s_k), r_m) \propto \exp(\kappa_k \eta_m(s_k)) \int_0^{\infty} \exp(-w_k \eta_m(s_k)^2/2) p(w_k|X^m(s_k) + r_m, 0) dw_k,$$

where $\kappa_k = (X^m(s_k) - r_m)/2$, $w_k$'s are independently distributed as $PG(X^m(s_k) + r_m, \eta_m(s_k))$, and $p(w_k|X^m(s_k) + r_m, 0)$ denotes the $PG(X^m(s_k) + r_m, 0)$ density. Then, for given $w_k$'s, defining $y_k = \frac{X^m(s_k) - r_m}{2w_k}$, we can write the posterior sampling distribution of $\boldsymbol{\beta}^{mm'}$ as

$$p(\boldsymbol{\beta}^{mm'}|.) \propto \pi(\boldsymbol{\beta}^{mm'}|.) \exp\left[-\frac{1}{2}(\mathbf{Y} - \mathbf{X}^{*m'}\boldsymbol{\beta}^{mm'})^T \Omega (\mathbf{Y} - \mathbf{X}^{*m'}\boldsymbol{\beta}^{mm'})\right], \ \Omega = diag(\mathbf{w}),$$

where $\mathbf{Y} = (y_1, \ldots, y_n)^T$ and $\mathbf{w} = (w_1, \ldots, w_n)^T$, and $\pi(\boldsymbol{\beta}^{mm'}|.)$ is from Eq. 4. Upon simplification, the posterior sampling step of $\boldsymbol{\beta}^{mm'}$ can be written as

$$p(\boldsymbol{\beta}^{mm'}|.) \propto \exp\left[-\frac{1}{2\sigma^2}(\boldsymbol{\beta}^{mm'} - M)^T \Sigma^{-1} (\boldsymbol{\beta}^{mm'} - M)\right],$$
$$\Sigma = \left(B + (\mathbf{X}^{*m'})^T \Omega \mathbf{X}^{*m'}\right)^{-1}, \ M = \Sigma\left((\mathbf{X}^{*m'})^T \Omega \mathbf{Y}\right).$$

Putting all the steps together, the Gibbs sampling steps are

1. For $k = 1, \ldots, n$, draw $w_k$ from $PG(X^m(s_k) + r_m, \eta_m(s_k))$, where $\eta_m(s_k) = \beta_0^{mm'}(s_k) + X^{m'}(s_k)\beta_1^{mm'}(s_k)$.

2. For $k = 1, \ldots, n$, define $y_k = \frac{X^m(s_k) - r_m}{2w_k}$.

3. Simulate $\boldsymbol{\beta}^{mm'}$ from $MVN(M, \Sigma)$, where

$$\Sigma = \left(B + (\mathbf{X}^{*m'})^T \Omega \mathbf{X}^{*m'}\right)^{-1}, \ M = \Sigma\left((\mathbf{X}^{*m'})^T \Omega \mathbf{Y}\right).$$

4. In case of the spatial fused lasso, consider steps 2 and 4 from Section 2.2.1, without the $\sigma^2$ term. In case of the spatial fused horseshoe, consider step 2 from Section 2.2.2, without the $\sigma^2$ term.

5. Update $r_m$ using a conjugate Gamma distribution and introducing latent terms $l_k$'s that follow the Chinese restaurant table (CRT) distribution, as described in Dadaneh et al. (2018) [13]

$$l_k \sim CRT(X^m(s_k), r_m),$$
$$r_m | l_1, \dots, l_n, . \sim Gamma(a + \sum_{k=1}^{n} l_k, b - \sum_{k=1}^{n} \psi_m(s_k)).$$

#### 2.4 Connection of the Gaussian model with bivariate spatial processes

In a univariate Gaussian process (GP)-based spatial model with an intercept and no covariates, it is typically assumed that $\mathbf{X}^m \sim MVN(\mu_m \mathbf{1}, \sigma_m^2 H + \sigma_E^2 I_n)$, where $H$ is a spatial covariance matrix with $H_{kk'} = \rho(||s_k - s_{k'}||, \phi)$, $\rho(.)$ is a known stationary correlation function, such as Matern [1], with hyperparameter $\phi$. $I_n$ is the $n \times n$ identity matrix. $\sigma_m^2$ and $\sigma_E^2$ are spatial and aspatial variances, respectively. In a bivariate (or multivariate) context, to model the dependency between the two variables $\mathbf{X}^m$ and $\mathbf{X}^{m'}$, usually a bivariate process with a separable Kronecker product-based covariance matrix is considered (see chapter 9 of Banerjee et al. (2008) [6],

$$(\mathbf{X}^m, \mathbf{X}^{m'})^T \sim MVN\left(\begin{bmatrix}\mu_m \mathbf{1}\\ \mu_{m'} \mathbf{1}\end{bmatrix}, \mathbf{\Sigma} = \underbrace{\begin{bmatrix}\sigma_m^2 & \nu\sigma_m\sigma_{m'}\\ \nu\sigma_m\sigma_{m'} & \sigma_{m'}^2\end{bmatrix}}_{T} \otimes H\right). \tag{6}$$

The autocorrelation of both variables has the form $corr(X^m(s_k), X^m(s_{k'})) = \rho(||s_k - s_{k'}||, \phi)$. While the parameter $\nu$ appears as an "at-location" correlation term since $corr(X^m(s_k), X^{m'}(s_k)) = \nu$, it also regulates the cross-correlation as $corr(X^m(s_k), X^{m'}(s_{k'})) = \nu\rho(||s_k - s_{k'}||, \phi)$ (between locations). Notice that the aspatial variance terms ($\sigma_E^2$) have been dropped to utilize the computational benefits associated with a Kronecker product structure, such as the ease of computing the inverse: $\mathbf{\Sigma}^{-1} = T^{-1} \otimes H^{-1}$. With appropriate priors on every parameter, a Bayesian model-fitting approach is straightforward.

To demonstrate how the Gaussian spatially varying coefficients (SVC) model (Eq. 1 from the main text) may alternatively be used to estimate $\nu$ and $\phi$, we focus on the conditional distribution [14] of $\mathbf{X}^m$,

$$\mathbf{X}^m|\mathbf{X}^{m'} \sim MVN(\tilde{\boldsymbol{\mu}}_m, \tilde{\boldsymbol{\Sigma}}_m),\ \ \tilde{\boldsymbol{\mu}}_m = \mu_m \mathbf{1} + \boldsymbol{\Sigma}_{12}\boldsymbol{\Sigma}_{22}^{-1}(\mathbf{X}^{m'} - \mu_{m'}\mathbf{1}),\ \ \tilde{\boldsymbol{\Sigma}}_m = \boldsymbol{\Sigma}_{11} - \boldsymbol{\Sigma}_{12}\boldsymbol{\Sigma}_{22}^{-1}\boldsymbol{\Sigma}_{21}, \tag{7}$$

where $\boldsymbol{\Sigma}_{11} = \sigma_m^2 H$, $\boldsymbol{\Sigma}_{12} = \nu\sigma_m\sigma_{m'}H$, and $\boldsymbol{\Sigma}_{22} = \sigma_{m'}^2 H$ are the respective blocks of the covariance matrix $\boldsymbol{\Sigma}$. The conditional mean and covariance matrix can be simplified as

$$\tilde{\boldsymbol{\mu}}_m = \mu_m \mathbf{1} + \nu\frac{\sigma_m}{\sigma_{m'}}(\mathbf{X}^{m'} - \mu_{m'}\mathbf{1}),\ \ \tilde{\boldsymbol{\Sigma}}_m = (1-\nu^2)\sigma_m^2 H.$$

Therefore, given $\mathbf{X}^{m'}$ and consistent estimators of $\mu_{m'}$, $\sigma_{m'}^2$ (such as sample moments obtained from marginal distribution) or simply upon standardizing $\mathbf{X}^{m'}$ by mean and SD, a linear regression model [14] becomes appealing to estimate $\nu$ and other parameters as

$$\mathbf{X}^m = \mu_m \mathbf{1} + \nu\frac{\sigma_m}{\sigma_{m'}}(\mathbf{X}^{m'} - \mu_{m'}\mathbf{1}) + \mathbf{w},\ \ \mathbf{w} \sim MVN\left(\mathbf{0}, \sigma_m^{*2} H\right),\ \ \sigma_m^{*2} = (1-\nu^2)\sigma_m^2. \tag{8}$$

where $\mathbf{w} = (w_1, \ldots, w_n)^T$ is a latent spatially varying random effect. In a Bayesian estimation, $\mathbf{w}$ is generally sampled at each MCMC step (could be marginalized otherwise). Thus, the correspondence with the SVC model (Eq. 1 from the main text) is found by letting $\beta_0(s_k) = \mu_m + w_k$ and $\beta_1(s_k) = \nu\sigma_m/\sigma_{m'} = \beta_1, \forall k \in \{1, \ldots, n\}$. With such a special case of the SVC model, one can assume a GP prior on $\boldsymbol{\beta}_0 = \mu_m\mathbf{1} + \mathbf{w}$ as $\boldsymbol{\beta}_0 \sim MVN(\mathbf{0}, \sigma_m^{*2}H)$, and pursue a Bayesian approach to estimate $\boldsymbol{\beta}_0, \beta_1, \sigma_m^*$, and $\phi$, from which the original parameters $\nu$ and $\sigma_m$ are easily recovered. $\mu_m$ is identifiable up to an additive constant. The extra noise term $\epsilon(s_k)$ in the SVC model (Eq. 1 from the main text) that we do not find in Eq. 8 might help in capturing any potential aspatial source of variation that is missed by the Kronecker structure (Eq. 6). Additional covariates such as $Z(s_k)$ may be added to the mean in Eq. 6 and the derivation remains similar.

A natural question arises: where does our model make simplifications compared to directly fitting the joint bivariate process from Eq. 6? Recall that the spatial covariance matrix $H$ is not fixed; instead, it is a function of the hyperparameter $\phi$. We have $\pi(\mathbf{X}^m, \mathbf{X}^{m'}) = \pi(\mathbf{X}^m|\mathbf{X}^{m'})\pi(\mathbf{X}^{m'})$. While the effect of disregarding $\pi(\mathbf{X}^{m'})$ might be minimal on the inference of $\nu$, we lose information on $\phi$, estimating it solely based on the spatial autocorrelation of $\mathbf{X}^m$. Moreover, it is unclear which of the two variables, $\mathbf{X}^{m'}$ or $\mathbf{X}^m$, should be conditioned on. If there is prior knowledge on the biological causal mechanism between the variables, one variable might be preferred over the other as the so-called outcome. For example, in the LR analysis, it might be more appropriate to treat receptor expression as a function of ligand expression. As a side note, the simple Pearson correlation is still an asymptotically unbiased estimator of $\nu$ but with a large variance due to the spatial autocorrelation present in both variables [15, 16, 17], which explains its inflated type 1 error in simulations. The spatially weighted cross-correlation measures, such as bivariate Moran's $I$ or Lee's $L$ statistic, are essentially Pearson correlation between spatially lagged variables, and therefore, produce inflated type 1 errors as well.

By treating $\beta_1(s)$ as a spatial process, our SVC framework leads to a more general bivariate spatial process of $(\mathbf{X}^m, \mathbf{X}^{m'})^T$, than Eq. 6. Conditional on the process $\beta_1(s)$, we can derive the covariance of the bivariate process as follows. In the Gaussian SVC model, let $\mathbf{X}^{m'} \sim MVN(\mu_{m'}\mathbf{1}, \sigma^2_{m'}H)$, $\boldsymbol{\beta}_0 \sim MVN(\mathbf{0}, \sigma^2_{m0}H)$ (independent of $\mathbf{X}^{m'}$), $\epsilon(s_k) = 0$, and $\boldsymbol{\beta}_1$ be a fixed vector. We have $cov(X^m(s_k), X^{m'}(s_k)) = cov(\beta_0(s_k) + \beta_1(s_k)X^{m'}(s_k), X^{m'}(s_k)) = \beta_1(s_k)\sigma^2_{m'}$ and at-location correlation becomes, $corr(X^m(s_k), X^{m'}(s_k)) = \frac{\beta_1(s_k)\sigma^2_{m'}}{\sqrt{(\sigma^2_{m0}+\beta_1^2(s_k)\sigma^2_{m'})\sigma^2_{m'}}} = \nu(s_k)$ (let) which is not a constant anymore. Autocorrelation takes the form, $corr(X^m(s_k), X^m(s_{k'})) = \frac{\rho(||s_k - s_{k'}||, \phi)(\beta_1(s_k)\beta_1(s_{k'})\sigma^2_{m'} + \sigma^2_{m0})}{\sqrt{(\sigma^2_{m0}+\beta_1^2(s_k)\sigma^2_{m'})(\sigma^2_{m0}+\beta_1^2(s_{k'})\sigma^2_{m'})}} = \rho(||s_k - s_{k'}||, \phi)(\nu(s_k)\nu(s_{k'}) + l(s_k)l(s_{k'}))$, $l(s_k) = \frac{\sigma^2_{m0}}{\sigma^2_{m0}+\beta_1^2(s_k)\sigma^2_{m'}}$, and cross-correlation becomes $corr(X^m(s_k), X^{m'}(s_{k'})) = \rho(||s_k - s_{k'}||, \phi)\nu(s_k)$ which is asymmetric in $(s_k, s_{k'})$. Using a multivariate notation,

$$\begin{bmatrix} \mathbf{X}^m \\ \mathbf{X}^{m'} \end{bmatrix} \sim MVN\left(\begin{bmatrix} \mu_{m'}\boldsymbol{\beta}_1 \\ \mu_{m'}\mathbf{1} \end{bmatrix}, \begin{bmatrix} \sigma^2_{m0}H + \sigma^2_{m'}D_{\boldsymbol{\beta}_1}HD^T_{\boldsymbol{\beta}_1} & \sigma^2_{m'}D_{\boldsymbol{\beta}_1}H \\ \sigma^2_{m'}HD_{\boldsymbol{\beta}_1} & \sigma^2_{m'}H \end{bmatrix}\right), D_{\boldsymbol{\beta}_1} = diag(\boldsymbol{\beta}_1). \quad (9)$$

When $\boldsymbol{\beta}_1 = \nu\frac{\sigma_m}{\sigma_{m'}}\mathbf{1}$ or $D_{\boldsymbol{\beta}_1} = \nu\frac{\sigma_m}{\sigma_{m'}}I_n$, and $\sigma^2_{m0} = (1-\nu^2)\sigma^2_m$, we have $cov(\mathbf{X}^m) = \sigma^2_m H$, $cov(\mathbf{X}^m, \mathbf{X}^{m'}) = \nu\sigma_m\sigma_{m'}H$, same as Eq. 6.

#### 2.5 Effect of underlying spatial graph on the spatial fused horseshoe prior

We investigate the assumption of independence between edge-wise differences of a spatially varying coefficient in the spatial fused horseshoe GMRF prior. Let us focus on the slope $\beta_1(s)$ for demonstration. Suppose $|E|$ denotes the total number of edges and $e_i = \beta_1(s_{k_i^1}) - \beta_1(s_{k_i^2})$ denotes the difference between coefficients corresponding to the locations $(s_{k_i^1}, s_{k_i^2})$ joined by the $i$-th edge. Under the assumption of independence, the prior distribution on $\mathbf{e} = [e_1, e_2, \ldots, e_{|E|}]^T$ can be written jointly as $\mathbf{e} \sim MVN(0, D),\ D = diag((\Lambda_{1i}^2 \tau_1^2 \sigma^2))_{i=1,\ldots,|E|}$. As discussed in the main text, $\mathbf{e}$ must also satisfy a set of implicit constraints corresponding to the cycles of $E$. For clarity, here $E$ represents a general spatial dependency graph, not the MST, which is acyclic. Let $\mathbf{A}$ be a $t \times |E|$ contrast matrix summarizing the constraints as $\mathbf{Ae} = 0$, where $1 \leq t \leq |E|$ denotes the number of cycles in $E$. Both Besag and Higdon (1999) [18] and Rue and Held (2005) [4] have heuristically argued, in the context of an intrinsic GMRF (IGMRF) prior, that such constraints do not alter the implied prior distribution on $\boldsymbol{\beta}_1$, or, in other words, $\mathbf{Ae} = 0$ does not need to be accounted for explicitly. For clarity, we provide a formal proof applicable to our context in the following theorem. In an IGMRF prior, $D$ is usually a scaled identity matrix, $\kappa I$ or more generally $\kappa\ diag((w_i))_{i=1,\ldots,|E|}$, where $w_i$'s are known constants. Therefore, conditional on $\Lambda_{1i}^2$'s, $\tau_1^2$, $\sigma^2$, we also have a special type of IGMRF prior.

**Theorem 1.** *The constraint* $\mathbf{Ae} = 0$ *does not influence the implied prior on* $\boldsymbol{\beta}_1$*,* $\pi(\boldsymbol{\beta}_1|.)$ *from Eq. 5 of the main text.*

*Proof.* Since $\pi(\boldsymbol{\beta}_1|.) = \pi(\mathbf{e})$ where $\mathbf{e} \sim MVN(0, D)$, it suffices to show that $\pi(\mathbf{e})$ does not change (up to a constant) under the constraint $\mathbf{Ae} = 0$, i.e., $\pi(\mathbf{e}|\mathbf{Ae} = 0) \propto \pi(\mathbf{e})$.

Using standard properties of an MVN distribution, conditional on $\mathbf{Ae}$ the distribution of $\mathbf{e}$ becomes a degenerate MVN distribution ($MVN_D$) [19] as

$$\begin{aligned} &\mathbf{e}|\mathbf{Ae} \sim MVN_D(0, P),\ \ P = D - D\mathbf{A}^T(\mathbf{A}D\mathbf{A}^T)^{-1}\mathbf{A}D, \\ &\log \pi(\mathbf{e}|\mathbf{Ae}) = -\frac{1}{2}\mathbf{e}^T P^+ \mathbf{e} + C \end{aligned} \tag{10}$$

where $P^+$ is the Moore Penrose inverse of the conditional covariance matrix $P$ and $C$ is a constant containing the pseudo-determinant. An $\mathbf{e}$, for which $\mathbf{Ae} = 0$, can be engineered using the conditioning by Kriging approach [4] which involves transforming an $\mathbf{e}$ simulated from the marginal distribution $MVN(0, D)$ as $\mathbf{e}^* = \mathbf{e} - D\mathbf{A}^T(\mathbf{A}D\mathbf{A}^T)^{-1}(\mathbf{Ae})$. It is easy to verify that $\mathbf{e}^*$ follows the same $MVN_D$ distribution from Eq. 10 and $\mathbf{Ae}^* = 0$. But, such a transformation is not explicitly required since $\log \pi(\mathbf{e}^*) = \log \pi(\mathbf{e}|\mathbf{Ae} = 0) = \log \pi(\mathbf{e}) + C^*$, for some constant $C^*$, as shown next. Letting $M = (I_n - D^{1/2}\mathbf{A}^T(\mathbf{A}D\mathbf{A}^T)^{-1}\mathbf{A}D^{1/2})$, we write $P = D^{1/2}MD^{1/2}$. $M$ is symmetric and idempotent since

$$\begin{aligned} M^2 &= (I_n - D^{1/2}\mathbf{A}^T(\mathbf{A}D\mathbf{A}^T)^{-1}\mathbf{A}D^{1/2})(I_n - D^{1/2}\mathbf{A}^T(\mathbf{A}D\mathbf{A}^T)^{-1}\mathbf{A}D^{1/2}) \\ &= I_n - 2D^{1/2}\mathbf{A}^T(\mathbf{A}D\mathbf{A}^T)^{-1}\mathbf{A}D^{1/2} + D^{1/2}\mathbf{A}^T(\mathbf{A}D\mathbf{A}^T)^{-1}\mathbf{A}D\mathbf{A}^T(\mathbf{A}D\mathbf{A}^T)^{-1}\mathbf{A}D^{1/2} \\ &= I_n - D^{1/2}\mathbf{A}^T(\mathbf{A}D\mathbf{A}^T)^{-1}\mathbf{A}D^{1/2} = M. \end{aligned}$$

Therefore, the Moore-Penrose inverse of $M$ is itself, $M^+ = M$. It implies that $P^+ = (D^{1/2}MD^{1/2})^+ = D^{-1/2}M^+D^{-1/2} = D^{-1/2}MD^{-1/2} = D^{-1} - \mathbf{A}^T(\mathbf{A}D\mathbf{A}^T)^{-1}\mathbf{A}$. Substituting $P^+$ in the log-likelihood from Eq. 10 and $\mathbf{Ae} = 0$,

$$\log \pi(\mathbf{e}|\mathbf{Ae} = 0) = -\frac{1}{2}\mathbf{e}^T(D^{-1} - \mathbf{A}^T(\mathbf{A}D\mathbf{A}^T)^{-1}\mathbf{A})\mathbf{e} + C = -\frac{1}{2}\mathbf{e}^T D^{-1}\mathbf{e} + C = \log \pi(\mathbf{e}) + C^*.$$

Thus, we complete the proof.

□

### 3 Convergence and run-time

In the majority of our analyses, we considered 5,000 MCMC iterations with 2,500 burn-in. However, the algorithm typically converges within the first 1,000 iterations. Therefore, for datasets with a large number of locations, a smaller number of MCMC iterations may be sufficient and can be safely employed when computational resources are limited. For the convergence diagnostics, we computed the Geweke statistic [20] for each $\beta_1^{mm'}(s_k)$, implemented in the *R* package *coda* [21], and investigated the trace plots of a few randomly chosen $\beta_1^{mm'}(s_k)$'s (Figs. 1 and 2). When either the variable $m$ or $m'$ is highly sparse ($> 75\%$ zeroes), imposing additional normal priors on $\beta_0^{mm'}(s_k)$'s and $\beta_1^{mm'}(s_k)$'s with a moderately large variance, such as $N(0, 5)$, drastically improves mixing and overall convergence performance. For one LR pair in the melanoma dataset, which has 292 spots, SpaceBF NB model takes 2 minutes (5000 MCMC iterations), on an Apple M1 Max-based MacBook pro with 10 cores and 64 GB RAM. SpaceBF Gaussian model is faster than the NB model. In the DCIS dataset, which has 5,548 spots, one run of the SpaceBF Gaussian model takes 24 minutes for 1000 MCMC iterations.

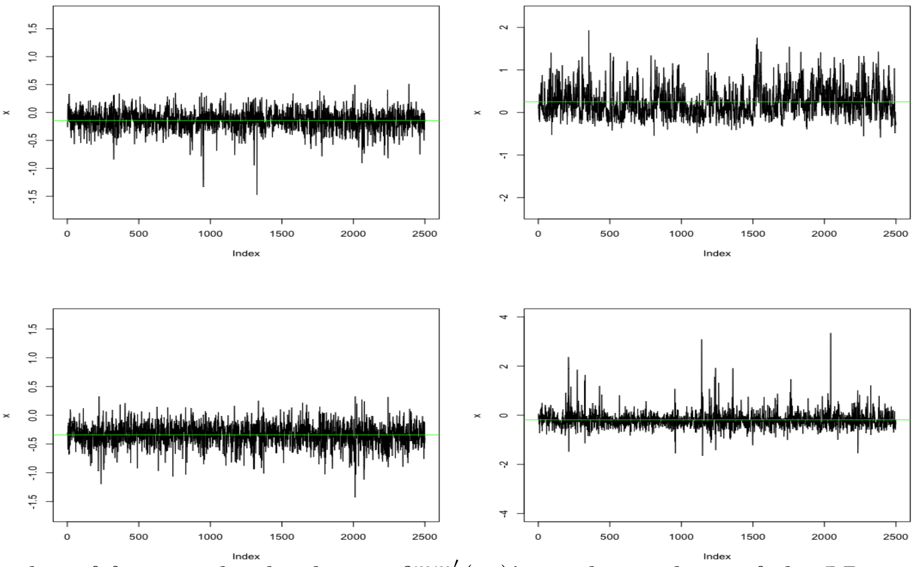


Figure 1: Trace plot of four randomly chosen $\beta_1^{mm'}(s_k)$'s in the analysis of the LR pair: (IGF2, IGF1R) from the melanoma dataset.

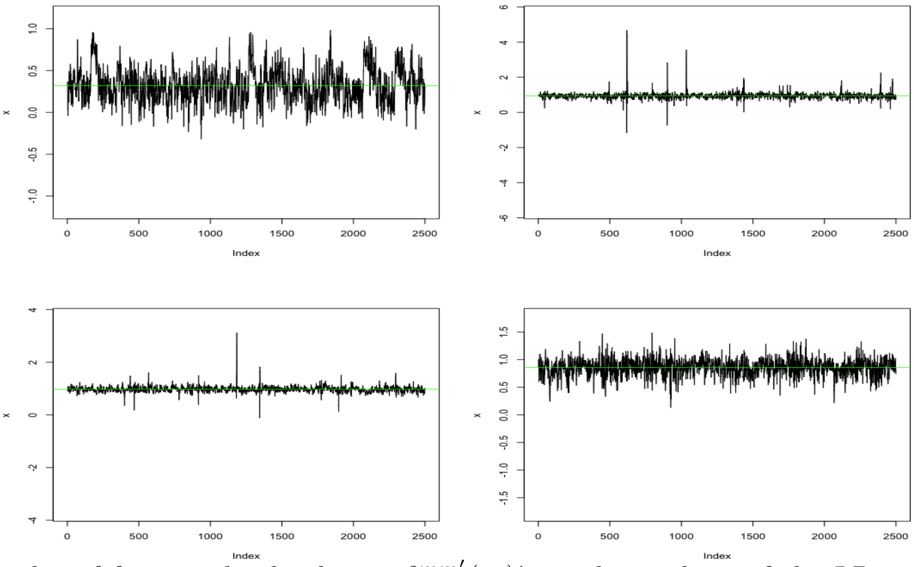


Figure 2: Trace plot of four randomly chosen $\beta_1^{mm'}(s_k)$'s in the analysis of the LR pair: (SPP1, CD44) from the melanoma dataset.
